# Intranucleolar Invasion of Cajal Body Remnants Suppresses Ribosomal Biogenesis in *cis* and Telomerase Functions in *trans*

**DOI:** 10.64898/2026.07.16.738943

**Authors:** Alexander D Morris, Julia Hoopman, Vidhyadhar Nandana, Cameron Guanxuan Lian, Emily Pochet, Maria Ruiz, Baihao Su, Andrey Efimov, Cynthia Myers, Yingda Tao, Luciano Saieva, Guoyu Lu, Erica Golemis, Livio Pellizzoni, Lu Chen

**Author notes:** These authors contributed equally (Alexander D Morris, Julia Hoopman).

## Abstract

Cellular processes are compartmentalized within immiscible heterotypic condensates, yet the functional consequences of losing their physical segregation remain unclear. Here, we show that genetic inactivation of the RNA chaperone SMN forces the aberrant intermixing of the two most prominent nuclear condensates: nucleolus and Cajal Body (CB). Upon SMN depletion, CB components invade the nucleolus and undergo reduced mobility and solubility consistent with a liquid-to-gel-like hardening transition. The CB-scaffold coilin aberrantly enriches at the nucleolar FC/DFC boundary and occupies rDNA chromatin, thereby locally suppressing rRNA production. Concurrently, this sequestration globally impairs coilin targeting to snRNA/snoRNA loci and limits telomerase access to telomeres, reducing telomeric synthesis. Crucially, genetic coilin depletion alone alleviates this mistargeting and rescues these functional impairments across condensates. Our findings reveal an inter-condensate rheostat model in which the loss of CB-nucleolar immiscibility is directly sensed, communicated, and executed by CB remnants, thereby proportionally coupling the functional outputs of otherwise distinct RNPs essential for splicing, translation, and genomic integrity.

## Introduction

Translation of mRNA to protein relies on the precise spatiotemporal coordination of ribonucleoprotein (RNP) complexes, a feat achieved by compartmentalizing the biogenesis of RNP machineries into membraneless organelles or liquid droplet-like condensates based on liquid-liquid phase separation (LLPS)^1^. To date, more than 40 distinct nuclear condensates have been curated in human cells^2^, many possessing unique molecular composition, specific viscoelastic properties, and dedicated biochemical functions. LLPS allows these to function as distinct organelles proximal to, but distinct from, their surrounding chromatin – itself the largest phase-separated cellular condensate^3^. While this “mixed-liquid” landscape redefines our view of nuclear organization, it presents a fundamental paradox: how do condensates enforce physical immiscibility while simultaneously communicating and coordinating their compartmentalized biochemical and genomic activities?

Originally identified as the most argyrophilic cellular structures in Ramón y Cajal’s silver staining^4^, the nucleolus and the CB (Cajal Body) are the principal nuclear sites for RNP biogenesis. Seminal studies of the nucleolus as a model condensate have shed light on intra-condensate dynamics^5,6^. As the nuclear site dedicated primarily to ribosome biogenesis, the nucleolus functions as a radially organized assembly line relaying rRNA across three immiscible layers. Nascent pre-rRNAs are initially transcribed from a core of rDNA chromatin or fibrillar center (FC), flow outwardly through the enveloping dense fibrillar component (DFC) for RNA processing, and eventually assemble with ribosomal proteins in an outermost shell, termed the granular component (GC). The directional FC-DFC-GC rRNA flow is associated with reduced viscosity and increased fluidity, with the FC/DFC interface resembling an entangled gel^5,6^.

The CB functions as a maturation^7^ hub for diverse RNP clients, including 1) snRNPs (**s**mall **n**uclear, i.e. U1, U2, U5), the core of the spliceosome^8^; 2) snoRNPs (**s**mall **n**ucle**o**lar, i.e. U3) and scaRNPs^9^ (**s**mall **Ca**jal Body), which guide the modification of rRNAs^10^ and snRNAs^11^, respectively; and 3) the telomerase^12^, a specialized scaRNP complex containing the scaRNA template (Telomerase RNA or TR)^13^ and a reverse transcriptase (TERT)^14^ that synthesizes telomere repeats at the ends of linear eukaryotic chromosomes^15^. Many CB client RNAs follow a salmon-like trafficking route: mature RNAs journey back home to CBs, which physically associate with the exact genomic loci where the nascent RNA counterparts were initially transcribed. In addition, CBs are known as “transcriptosomes”, concentrating the transcription apparatus from all 3 RNA polymerases^16^, including mediators^17^ and elongation factors^18^. Similar to the nucleolus, CBs exhibit liquid-like properties and a confined boundary that depends on ongoing transcription^19^ and snRNPs^20^.

Reflecting their common roles in RNP biogenesis, the nucleolus and CBs share a common cellular pool of major RNA-modifying machineries, including the dyskerin tetramer (for Box H/ACA-guided pseudo-Uridylation) and the fibrillarin tetramer (for Box C/D-guided 2’-O-methylation). In addition, CBs, once noted as the “nucleolar accessory body”, are frequently physically tethered to the nucleolar periphery^21^. Nevertheless, despite their intimate relationship^22,23^, CBs persist as an evolutionarily conserved, physically immiscible condensate that is functionally distinct from the nucleolus.

Notably, the CB-nucleolus immiscibility barrier is not immutable, but lost in a number of pathogenic conditions. Seminal ultrastructure^24^ and histology studies^25–27^ of neurodegenerative models, including Purkinje cell degeneration (PCD)^27^, amyotrophic lateral sclerosis (ALS)^28^, and spinal-muscular atrophy (SMA)^26,29^, have revealed striking phenotypes of CB reorganization, including the intranucleolar presence of the CB-defining scaffold coilin^16^, which is typically restricted to a localization within the CB^21,23^. Whether this loss of immiscibility is a driver or passenger effect in disease pathogenesis remains poorly understood.

Coilin is a potent nucleator for other CB-enriched client proteins, including TCAB1/WDR79^30,31^ and SMN^32^ – key biogenesis factors for scaRNPs^33^ and snRNPs^34^, respectively. Typical of an RNP condensate scaffold, coilin contains an N-terminal self-association domain, a central intrinsically disordered region (IDR), and an RG (Arginine-Glycine)-rich motif, contributing to its capacity to oligomerize^35^, its susceptibility to regulation by post-translational modifications^36^ such as sDMA (Symmetric Di-Methyl-Arginine), and a general affinity for RNA substrates^37,38^, respectively. In addition, coilin has been demonstrated by crosslinking-based proximity assays to occupy specific genomic loci encoding snRNAs, snoRNAs, and histone genes^38–40^, and is implicated in genome organization^39,41^.

Intriguingly, inherited mutations in the CB clients TCAB1 and SMN cause the human degenerative disorders *dyskeratosis congenita* (DC)^42^ and SMA^32^, respectively. Functionally, TCAB1 binds to the scaRNA CAB box, which is a defining feature of this class of small RNAs, and retains scaRNAs within the CB, diverting them from the nucleolar trafficking route taken by TCAB1-free snoRNAs^12,30,31^. In addition, TCAB1 is required for the activity and telomeric targeting of the telomerase scaRNP^43,44^, suggesting a molecular explanation for the premature aging associated with DC. SMN chaperones the assembly of Sm rings to pre-snRNAs^34^, and together with TGS1^45^, constitutes a bipartite requirement that licenses the nuclear re-import of snRNPs to the CB^46^. While the neurodegenerative disease SMA is associated with pleiotropic molecular phenotypes, the splicing dysfunction caused by insufficient snRNP dosage has classically been recognized as the primary driver for motor neuron loss^47^. However, the fact that SMN dysfunction is linked to loss of immiscibility also or alternatively suggests a potential driver role for disruption of LLPS in disease pathogenesis.

While intranucleolar coilin has often been applied as a useful histological stress marker, the functional consequence of this displacement has remained largely obscure. Among related questions, it has not been clear whether other proteins usually resident in the CB travel along with the nucleolar coilin and whether mislocalized CB scaffold actively reshape the NU. Whether CB RNPs still function properly under conditions that induce displacement is unknown. Critically, the biophysical properties of a CB condensate that breaches its native phase boundary have been undefined.

Here, we model the biological impacts of inter-condensate crosstalk by using SMN depletion to enforce stable, population-wide intermixing between the CB and nucleolus. This allows us to address critical unanswered questions regarding the mechanisms of inter-condensate crosstalk. Focusing on the definitive CB scaffold coilin, we use live cell imaging to show that directed migration of coilin into the nucleolus is accompanied by multiple CB remnants. CB components within the invaded nucleolus undergo a liquid-to-gel transition, with intra-nucleolar coilin failing to bind its native genomic targets while acquiring *de novo* occupancy for rDNA chromatin at the FC/DFC interface. We further show that coilin is indispensable in the intranucleolar liquid-to-gel phase hardening of CB remnants, a process that simultaneously compromises rRNA transcription *in cis* and telomerase functions *in trans*. Together, these insights emphasize the role of coilin in maintenance of LLPS, and the barriers that enforce or allow inter-condensate communication.

## RESULT

### Coilin-directed CB nucleolar invasion upon SMN depletion

To trigger loss of CB-nucleolus immiscibility, we used a stable dox-inducible shSMN cell line to deplete SMN. Within 3 days after dox treatment, we observed rapid depletion of SMN protein (Fig.1A, Extended Data Fig.1B), SMN mRNA (Extended Data Fig.1A), and multiple subunits of the SMN-associated GEMIN complex (Extended Data Fig.1B). Although SMN was depleted by Day 3, the population doubling rate remained stable until Day 5 (Fig. 1B), after which the growth rate slowed. Concurrently, beginning from Day 5, we detected intranucleolar accumulation of CB components, including the CB scaffold coilin, the scaRNA TR (Fig.1C), the scaRNP chaperone TCAB1 (Extended Data Fig.1C), and the CB snRNA processing enzyme TOE1 (Extended Data Fig.1D). This phenotype increased progressively upon prolonged SMN depletion (Extended Data Fig.1E), resulting in a gradually increased colocalization of coilin and the nucleolus (Fig.1C-D; delineated by nucleolar markers and characteristic DAPI-low regions^48^).

**Fig.1.**
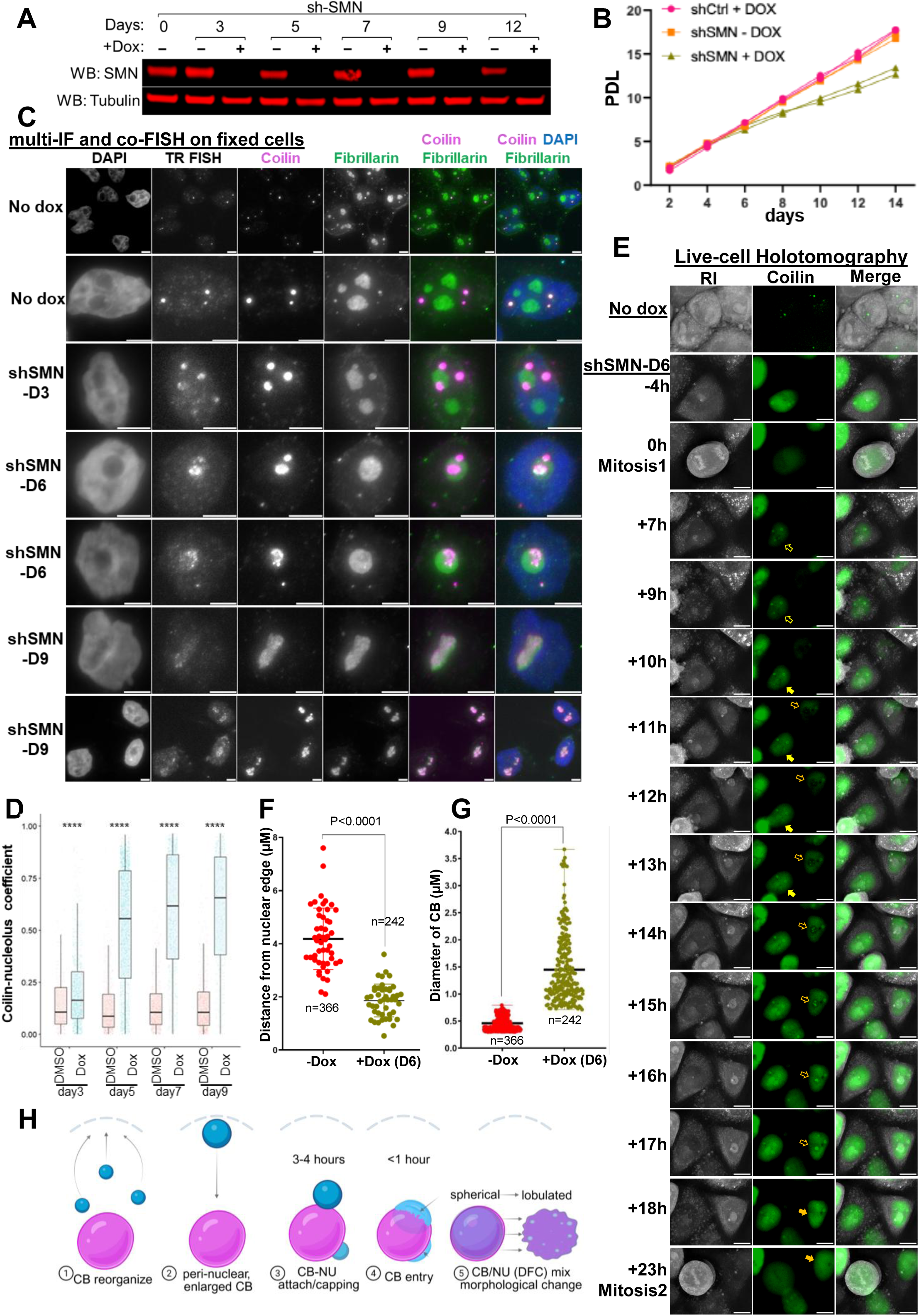
The Cajal Body undergoes nucleolar invasion upon SMN depletion. (A) WB of SMN protein accumulation in stable HeLa cells expressing Dox-inducible shRNA targeting human SMN genes (sh-SMN). (B) PDL growth curve, cumulative Population Doubling Level shown in HeLa cells expressing control shRNA or shSMN +/- Dox at indicated days. (C) Co-IF of the HeLa shSMN stable cell line +/- Dox at indicated days. Coilin, the CB marker in pink; Fibrillarin, a marker of the nucleolar DFC in large. Scale bars, 3 µm. (D) Coilin-nucleolus colocalization during the time course of Dox treatment. Object-based colocalization between segmented CBs and NUs were quantitated using CellProfiler. Kruskal-Wallis test considering the non-normal distributions was used with p≤ 0.0001, n= 8,439 (E) Live cell holo-tomography imaging frames. shSMN HeLa cell line stably expressing EGFP-tagged coilin, after 6 days of Dox treatment. Two consecutive mitoses shown. Two daughter cells from the first mitosis were marked with yellow or orange arrows, respectively. Filled arrows indicate the occurrence of coilin’s nucleolar invasion. Scale bar, 5 µm. (F) Diameter of Coilin-positive structures upon 6 days of Dox treatment. n=366 for - Dox condition, while n=242 for + dox condition. (G) Distance of Coilin-positive structure from nuclear edge. N=366 for -Dox, while n=242 for +dox. (H) An illustration to describe the major intermediate steps during and after CB’s nucleolar invasion.

To temporally resolve the intermediate steps leading to CB’s nucleolar invasion, we performed live-cell holo-tomography in an EGFP-coilin stable cell line following the longitudinal depletion of SMN (Fig.1E, Extended Data Fig.1E). Supporting a loss of the canonical CBs, the nucleoplasmic EGFP-coilin signal increased (vs. the No Dox, Fig.1E), where we detected coilin-positive spherical shells near the nuclear periphery (Fig.1F). These coilin shells are ∼ 3-fold larger than the typical CB (Fig.1G), and encompass an inner core positively for TCAB1 (CB) as well as nucleolar proteins from all three phases, including nucleolin (from the GC; Extended Data Fig.2H), fibrillarin (from the DFC; Fig.1C, Extended Data Fig.1E), in addition to the RNA Pol-I subunit RPA194, and the rDNA transcription factor UBF1 (from the FC; Extended Data Fig.2A). Frequently, these coilin shells translocate inwardly to dock on the nucleolar surface (Extended Data Fig.1F), transforming subsequently into half-dome-shaped peri-nucleolar caps (Extended Data Fig.1E-F). In addition, coilin colocalizes with low-RI (Refractive Index) areas within the high-RI backdrop of nucleoli, exhibiting dynamic liquid-like fission and fusion behavior for up to 6 hours (Fig.1E and Extended Data Fig.1E). Within 1 hour, coilin-positive caps and the low-RI structure simultaneously disappear, coinciding with the spread of coilin intranucleolarly (Fig.1E, H). Morphologically, the resulting CB-nucleolar mixture underwent a spherical-to-lobulated morphological change, increasing its eccentricity (Fig. 1H, Extended Data Fig.1G). While the sequential shell-to-cap-to-invasion transition can unfold over several hours within a single interphase (Fig.1E and 1H), prolonged SMN depletion accelerates this process, resulting in a predominantly intranucleolar coilin phenotype after several days of dox-treatment (Fig.1C, Extended Data Fig.1E).

**Fig.2.**
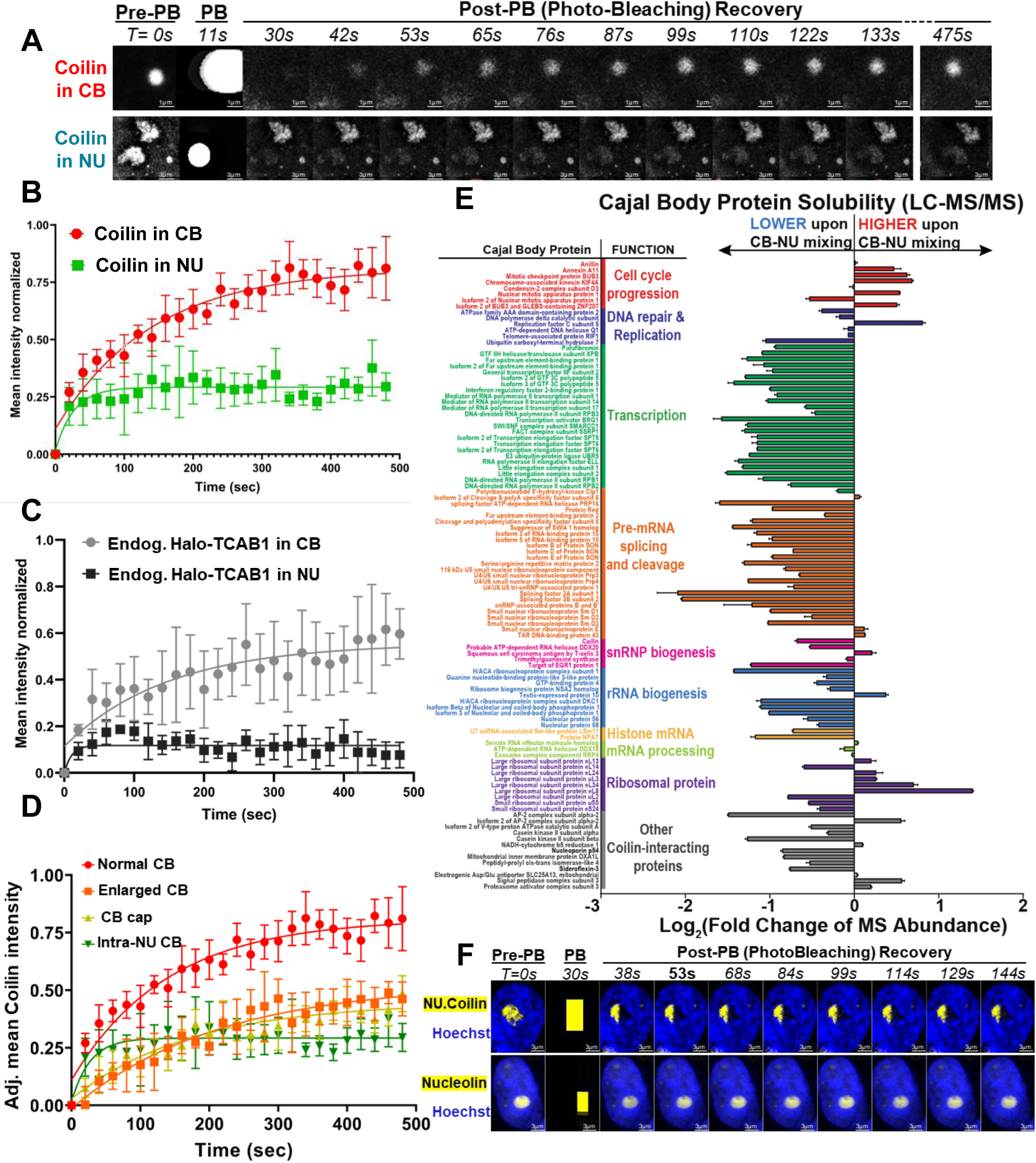
Intranucleolar Coilin shows decreased mobility and solubility. (A) FRAP on Coilin-positive structures, either the conventional CB or the intranucleolar CB, from 10-day Dox-treated sh-SMN cells transfected with EGFP-Coilin. PB, photobleaching; the PB frame describes the area of photobleaching. A series of post-PB frames is shown. (B) Adjusted mean EGFP intensity from Fig. 2A, sum-projected Z-stacks, were plotted over the time course (n=10, fit with single exponential function). Scale bar, 95% confidence interval. (C) FRAP dynamic of Halo-TCAB1 from shSMN cells carrying an endogenously knocked-in Halo tag at the ATG of the TCAB1 gene. TCAB1 in CB, no dox treated, n=9; TCAB1 in NU, dox-treated for 11 days, n=6. (D) FRAP dynamic of EGFP-Coilin as in Fig.2A. 3 types of Coilin-positive structures were compared from Dox-treated shSMN cells. n= 8, 6, 6, respectively. (E) LC-MS/MS abundance of CB proteins with functional annotations, per Escayola et al, in a post-NP40 chromatin fraction solubilized by the RIPA detergent and DNaseI. Abundance values were normalized using the MaxLFQ algorithm and performed in duplicates, with SD shown. (F) Partial nucleolar FRAP in dox-treated (10 days) shSMN cells transfected with either EGFP-tagged Coilin or Nucleolin. Approximately half of the NU is photobleached, and the recovery frames are shown.

Thus far, we have shown that SMN-depleted CB remnants reorganize and adopt a specific trafficking pathway leading to nucleolar invasion. To determine whether coilin maintains its scaffolding role within CB remnants, we generated isogenic coilin-knockout cell clones, validated by WB and allelic sequencing (Extended Data Fig.1J-K). We found that intranucleolar invasion of TCAB1 strictly depends on the presence of coilin (Extended Data Fig.1H-I). Notably, coilin-deficient nucleoli still display the distinct low-RI structure, suggesting it is not a CB remnant (data not shown). The FC, being the least dense nucleolar phase^49^, is known to translocate from the nucleolar core to surface upon rRNA transcription stalling^50^. Therefore, we propose that the exposed FC may serve as a direct landing site during nucleolar stress, recruiting a CB-nucleolus hybrid intermediate from the nuclear periphery. Indeed, leveraging the nuclear periphery to manage rRNA biogenesis mirrors the evolutionary adaptation in the protozoan Tetrahymena, whose numerous nucleoli are arranged on the nuclear periphery^51^.

### Intranucleolar CB remnants show decreased mobility and solubility

Liquid-like condensates exhibit spherical morphology and dynamic behaviors with their surrounding environment^52^. The drastic reorganization of CB components from a spherical CB into a lobulated CB-nucleolar hybrid prompted us to test whether there was an underlying phase transition. We performed FRAP (Fluorescence Recovery After Photobleaching) in cells stably expressing either GFP-coilin or endogenously tagged Halo-TCAB1^43^. Within spherical CBs, coilin and TCAB1 exchanged with the nucleoplasm fairly freely (t_1/2_<100s), and the majority of these two proteins resided in the mobile fraction (Fig. 2A-C). In contrast, nucleolar internalized coilin showed markedly retarded fluorescence recovery after photo-bleaching (Fig. 2A-B). Consistently, the fraction of coilin and TCAB1 molecules in their respective mobile phase was reduced by 70% (Fig. 2B-C). Strikingly, a reduced efficiency of FRAP for coilin (Fig. 2D) was detected in all 3 intermediate CB structures (Extended Data Fig.1D) upon SMN depletion. In a complementary assay, LC/MS analyses of DNaseI-solubilized chromatin fractions suggested a systematic reduction in CB protein^22^ solubility upon CB-nucleolar mixing (Fig. 2E).

Consistent with a lack of intranucleolar mobility, in a partially photobleached nucleolus, GFP-coilin showed no detectable FRAP (Fig. 2F), suggesting a loss of liquid-like behavior. In contrast, GFP-nucleolin, which typically localizes to the liquid-like GC component, recovered fully and comparably to the level of the unbleached half of the nucleolus, within 8 seconds of photobleaching (Fig. 2F). Beyond their differences in mobility, the CB and the GC behave as strictly immiscible phases before, during, and after nucleolar invasion (Extended Data Fig.2A), whereas the invaded CB and the FC are readily miscible (Extended Data Fig.2B). Furthermore, we show that the GC layer, marked by NPM1, undergoes localized thinning precisely at the CB docking sites on the nucleolar periphery (Extended Data Fig.2A). Consistent with a localized breakdown of GC integrity, we observed a corresponding depletion of peri-nucleolar heterochromatin specifically at these docking sites (Extended Data Fig.2C-D). This aligns with previous studies showing that the core GC scaffold^49^, NPM1, is essential for maintaining the perinucleolar heterochromatin architecture^53^. Together, these imaging data suggest that the process of CB nucleolar invasion may be facilitated by the localized breaching of an otherwise immiscible GC barrier and its associated heterochromatin.

As an orthogonal means to induce CB-nucleolar mixing, we treated cells with CX-5461, a G-quadruplex (G4) stabilizer and potent RNA polymerase I (Pol I) inhibitor known to compromise nucleolar integrity by inhibiting rRNA transcription. We observed intermixing between coilin and fibrillarin (DFC), while mutual exclusivity between coilin and NPM1 (GC) remained (Extended Data Fig.2F-G). Complementary live cell imaging using cells expressing Halo-tagged TCAB1 and GFP-tagged nucleolin also indicated that, despite the loss of nucleolar integrity, CB and GC remain immiscible. While homotypical fusions between CB-to-CB and GC-to-GC occurred frequently, endogenous Halo-TCAB1 at the CB-GC interface exhibited typical liquid-like behaviors^54^. These included dynamic protrusions (fingering), interfacial fluctuations (ruffling), and surface spreading (wetting) along the nucleolar periphery (Extended Data Fig.2H).

Together, these results demonstrate that the dynamic and mobile nature of CB components is diminished upon their invasion into the nucleolus. The additional loss of droplet-like spherical character and solubility of the resultant CB/nucleolar mixture implies an underlying liquid-to-gel transition of intranucleolar CB remnants.

### Intranucleolar coilin binds rDNA in a restricted peri-FC zone

We reasoned that the liquid-to-gel transition of the intranucleolar CB following nucleolar invasion could be related to the acquisition of *de novo* interactions with unintended substrates within the non-native nucleolar environment. Confocal imaging pinpointed the intranucleolar location of coilin to the periphery of the FC (indicated by signal localization adjacent to the RNA pol I subunit RPA194, Fig. 3A, Extended Data Fig.3A). This colocalization between coilin and RPA194 progressively increased for more than 3 days following SMN depletion (Fig. 3B). Strikingly, a 3D reconstruction of the confocal images showed that coilin formed a tubular structure that connected most, if not all, FCs (Fig. 3C). Unexpectedly, rather than mixing throughout the nucleolus, coilin molecules were spatially confined within a subset of the FC/DFC interface (Fig.3C, Extended Data Fig.3B) and remained notably absent from the bulk of nucleolar volume marked by fibrillarin (Fig.3C). While all detected RPA194-positive FCs nucleated fibrillarin-positive DFCs, the percentage of RPA194 in contact with coilin was consistently lower than FC-DFC across cells (Fig. 3E, Extended Data Fig.3B). At these contact sites, coilin exhibited liquid-like fusion and surface wetting against its FC boundary (Fig. 3D). Together, these observations indicate that the nucleolar invasion of CB remnants is not a process of random diffusion, nor a complete mixing with DFC, but rather a targeted accumulation at the FC/DFC interface.

**Fig.3.**
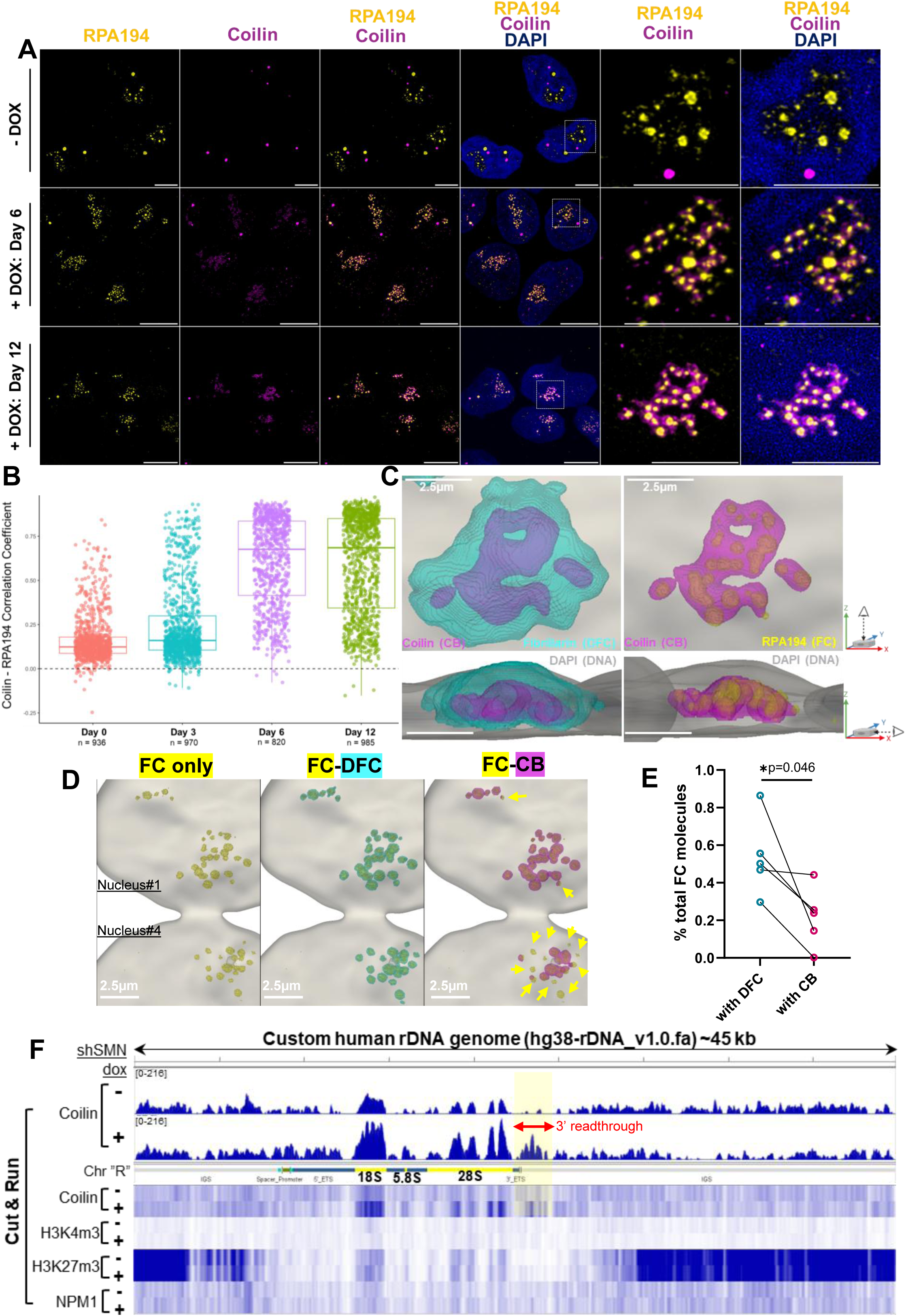
Intranucleolar Coilin binds rDNA at a restricted peri-FC zone. (A) Confocal microscopy of co-IF from HeLa shSMN cells treated with Dox as indicated. FC (RPA194, yellow), DFC (fibrillarin, cyan), CB (coilin, magenta), and DNA (DAPI, blue). Maximum Intensity Projections of Z-stacks are shown. Scale bar, 5 µm. (B) The degree of coilin-RPA194 colocalization during the course of Dox treatment. (C) 3D reconstitution of the confocal images in Fig.3A. Top-down view of the cell on the top row, side view on the bottom row. FC (RPA194, yellow), DFC (fibrillarin, cyan), CB (coilin, magenta), and DNA (DAPI, grey). (D) 3D reconstitution of the two nuclei #1 and #4 quantitated in Extended Data Fig.3B, showing overlays among FC (RPA194, yellow), DFC (fibrillarin, cyan), CB (coilin, magenta), and DNA (DAPI, grey). Yellow arrows denote the FC regions either partially or completely devoid of the coating by CB, while fully enveloped by DFC. (E) Linked percentage of the FC molecules (both surface and core) in contact with DFC or CB. Each linked data point sums up all molecular contacts within a single cell (n=5). P=0.046 (F) CUT&RUN reads of Coilin, H3K4me3, H3K27me3, and NPM1 mapped to a customized single-copy human rDNA genome sequence. sh-SMN cells treated with +/- dox for 10 days. Specific gain of coilin footprints at the downstream of the 3’ ETS in the 10-day Dox-treated sample is shaded in yellow.

The FC/DFC interface forms a specialized nucleolar zone highly enriched in actively transcribing rDNA repeats^6^. If invaded CB remnants stably occupy this zone, it should predictably result in *de novo* binding between coilin and the rDNA chromatin. To map coilin’s genomic occupancy of rDNA prior to and following nucleolar invasion, we performed genome-wide CUT&RUN profiling of endogenous coilin (Fig. 3F). Importantly, global coilin expression was not affected by SMN depletion (Extended Data Fig.3C), providing a controlled system to interrogate the gain or loss of SMN-dependent coilin footprints.

Bioinformatically mapping the entirety of coilin CUT&RUN reads to a single-copy consensus rDNA genome^55^ allows a direct quantitative comparison of coilin footprints. We found that SMN depletion significantly elevated coilin binding across the rDNA chromatin (Fig. 3F). This expanded coilin footprint spread broadly across the rRNA consensus, including the 5’ external transcribed sequence (ETS), 18S/5.8S/28S-encoding sequences, and, unexpectedly, a 1.5 kb noncoding region downstream of the 3’ ETS (Fig. 3F). As negative controls, the CB-immiscible GC marker NPM1 showed no specific footprint on rDNA genes. Conversely, the repressive peri-nucleolar heterochromatin mark H3K27me3^56^ was robustly distributed throughout the Intergenic Spacers (IGSs) regardless of SMN status (Fig. 3F). Together, these genomic profiles confirm our spatial observations, demonstrating that the nucleolar invasion of coilin culminates in its direct physical anchoring to the active rDNA transcription units.

### Intranucleolar coilin impedes rRNA transcription

The observed gain of coilin occupancy at rDNA repeats and the liquid-to-gel hardening of intranucleolar CB remnants could affect rRNA transcription. Using a ^32^P-orthophosphate pulse-chase assay^57^, we monitored the kinetics of nascent rRNA synthesis and maturation following SMN depletion (Fig. 4A, B). Strikingly, we found a 50% reduction in the production of not only the pre-rRNA species (47/45S) but also all labeled rRNA species (Fig. 4B-C), whereas the corresponding steady-state rRNAs remained unaffected (Fig. 4B). In addition, the maturation rates of both 28S and 18S rRNAs were reduced by ∼50% (Fig.4D). One explanation for such a decline could be defective snoRNA supply induced by SMN depletion, since snoRNAs are required for pre-rRNA cleavage^5^. However, we did not observe an accumulation defect of U3 snoRNA or TR scaRNA (Extended Data Fig.4A). As an orthogonal approach to labeling nascent rRNA, we pulse-chased nascent RNAs with 4-thiol-uridine (4sU), followed by biotinylation and purification of nascent RNA transcripts (Extended Data Fig.4B). Consistently, the SMN-depleted cells exhibited a reduced level of 4SU/Biotin in the bulk nascent RNAs at 1 and 8 hours after the initial pulse (Extended Data Fig.4C). We confirmed that this reduction is primarily driven by reduced rRNA species using RNA Tapestation electrophoresis (Extended Data Fig.4D). Together, these orthogonal nascent rRNA kinetics reveal that the nucleolar invasion of coilin impairs *de novo* rRNA synthesis independent of the steady-state ribosome pool.

**Fig.4.**
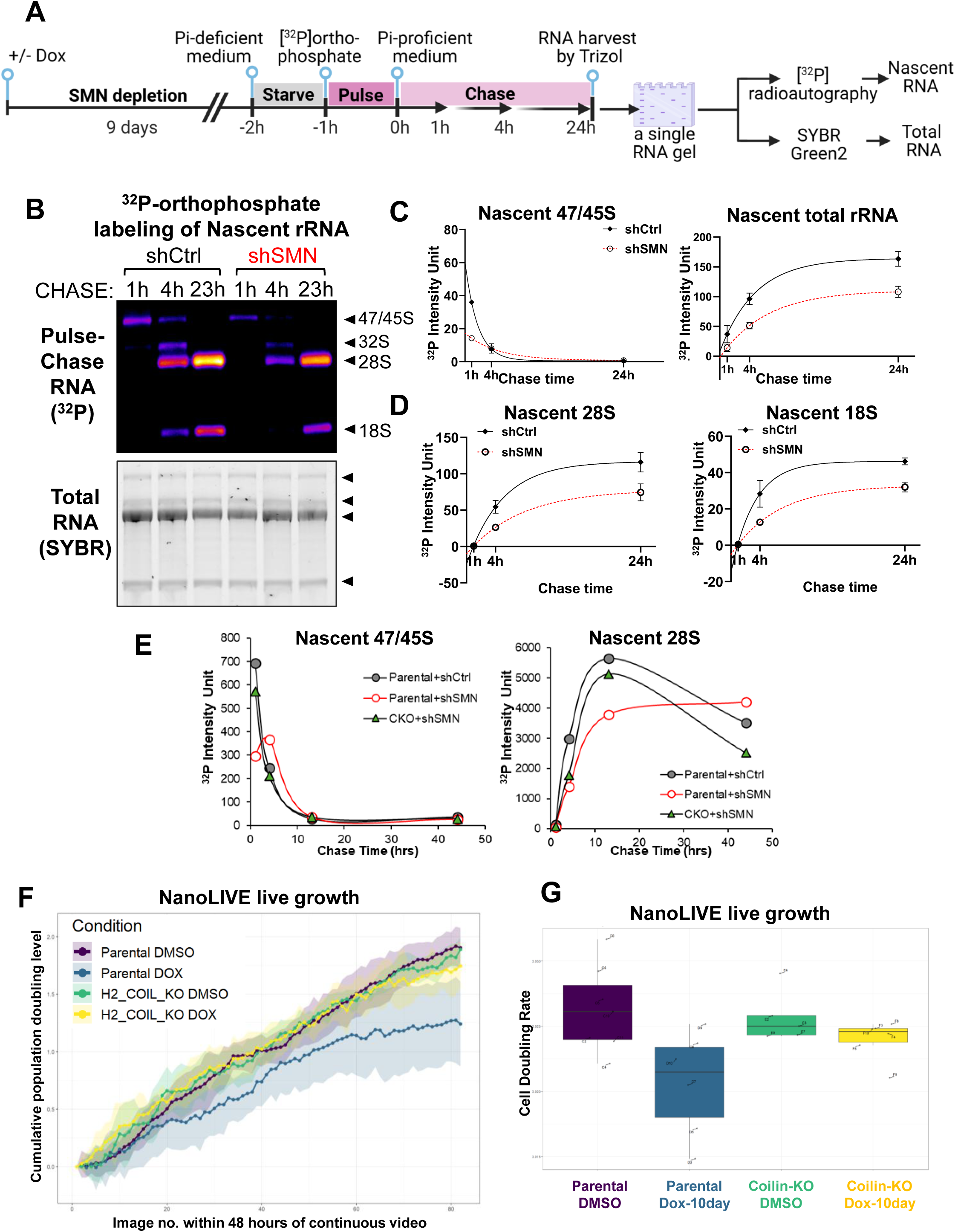
Intranucleolar Coilin impedes rRNA transcription. (A) Quantitation of nascent rRNA production by pulse and chase labeling shSMN cells (Day-9 Dox-treated or mock) with ^32^P-orthophosphate. (B) Pulse-chase labeling of nascent rRNA production using ^32^P-orthophosphate, in either shCtrl or shSMN expressing cells treated with dox for 10 days. (Top) the agarose-MOP gel is exposed by autoradiography to visualize labeled rRNA species at the indicated chase time; (Bottom) the exact same gel was stained with SYBR-Green-2 dye for normalization purposes. Dark triangles indicate the predicted rRNA species. Representative images from two replicates are shown. (C) The quantitation of nascent rRNA from the B panel. Arbitrary raw unit counts from ^32^P-orthophosphate autoradiography are shown for the nascent 47/45S band and for total rRNA bands, combining all detectable rRNA species including 47/45S, 32S, 28S, and 18S. (D) The quantitation of the nascent 28S or 18S band from B panel. (E) An isogenic Coilin-KO clone (H2) was treated with SMN depletion, and rRNA production measured by the rRNA pulse-chase assay using ^32^P-orthophosphate. The processing of nascent 47/45S and the production of 28S are quantified with coilin-proficient parental cells. (F) Nanolive live imaging recording of continuous cell proliferation. X-axis, image number evenly distributed along a 48-hour imaging window. The Y-axis, real-time cumulative population doubling level, extrapolated from 24 individual wells from a 96-well plate. Four conditions, parental vs. H2 clone of the Coilin-KO, with DMSO or DOX for 10 days. (G) Box plot of the cell doubling rate from the same experiment as in F. Growth rate from individual 96-well plate is labeled.

Having established that coilin retains its scaffolding properties within the nucleolus (Extended Data Fig.1H-I), we asked whether it is directly responsible for the attenuated rRNA synthesis. Indeed, the targeted knockout of *COILIN* (Extended Data Fig.1J-K) fully restored the production kinetics of 47S pre-rRNA and the subsequent maturation of the 28S rRNA despite the prolonged SMN depletion for 10 days (Fig. 4E, Extended Data Fig.4E). This suggests that the aberrant accumulation of intranucleolar coilin is the primary driver of the rRNA biogenesis defect. Importantly, *COILIN* knockout also restored normal population doubling rates (Fig. 4G, 1B), mechanistically linking the suppression of rRNA production by invading coilin to the resulting proliferative impairment.

### Genomic targeting of coilin is compromised by CB-nucleolar mixing

Prior ChIP-seq profiling^38^ showed that coilin occupies intronless genes, including those loci encoding snRNAs, snoRNAs, and replicative histones - the core histone clusters transcribed strictly during S phase. To determine whether coilin retains these interactions in intranucleolar CB, we profiled the genome-wide occupancy of coilin using CUT&RUN. Among the GENCODE-annotated snRNAs and snoRNAs, we detected coilin peaks mapped to only ∼1% of these loci (Fig.5A), many of which are snRNA repeats located on the same chromosome (Extended Data Fig.5A). In SMN-proficient cells, coilin peaks at the genomic regions downstream of the transcription end site (TES), extending up to 1 kb beyond the annotated 3’ ends (Fig.5B). Interestingly, the 3’ bias of coilin targeting is reminiscent of the expanded coilin footprint that we observed downstream of the rDNA 3’ ETS (Fig. 3F). However, in SMN-depleted cells, the coilin footprint at these native sn/snoRNA targets is markedly diminished. Together, these data suggest that while coilin retains a targeting preference for the 3’ readthrough regions, its access to native genomic targets is abolished upon nucleolar invasion.

**Fig.5.**
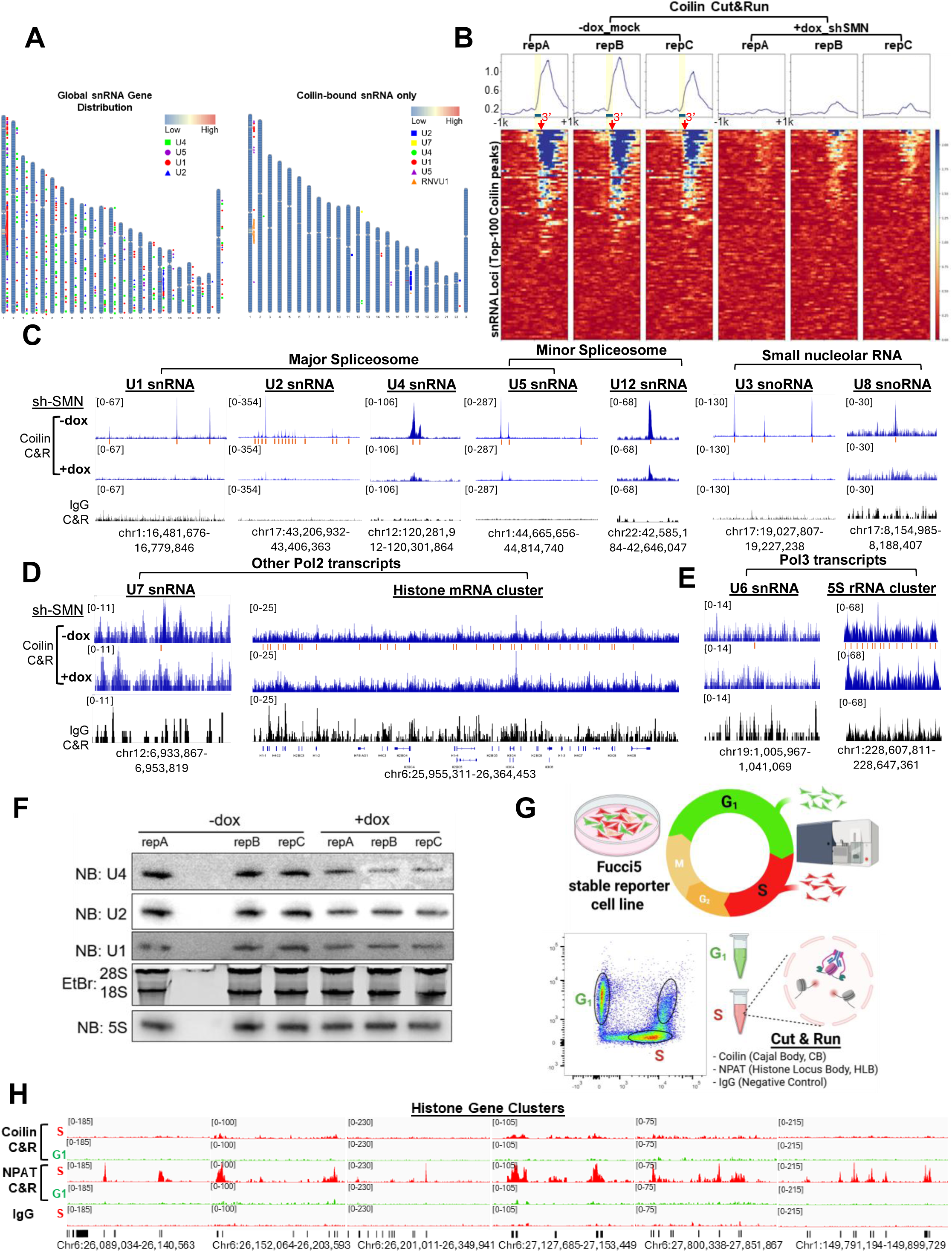
Coilin’s genomic targeting is compromised upon CB-NU fusion. (A) Ideograms on the left showing the genomic distribution of annotated snRNA gene loci for U4, U5, U1 and U2, and on the right, only those coilin-bound snRNAs per CUT&RUN were labeled. The relative CUT&RUN signal intensity is represented by a blue (low) to red (high) color scale. (B) Heatmap showing the top one hundred Coilin CUT&RUN peaks by centering the Transcription Start Site (TSS) within a 2kb genomic window; +/- dox-induced shSMN for 10 days; 3 replicates shown. The summary plots are shown on top, with the snRNA-encoding gene body labeled as a Teal bar and shaded in yellow. The genomic position for the annotated 3’ end of the snRNA gene body is indicated by a red arrow. (C-E) IGV browser views of the coilin CUT&RUN peaks at indicated genomic regions, shSMN cells +/- dox for 10 days; IgG was included to control for antibody specificity. Individual coding sequences of indicated transcripts are denoted underneath the track by red bars. (F) Northern Blotting of steady-state snRNA accumulation in shSMN cells, +/- 10-day of Dox-treatment shown. 3 replicates are included. (G) Cell cycle-specific CUT&RUN with shSMN cells stably expressing a Fucci5 reporter. G1 (positive for green fluorescence) or S cells (positive for red fluorescence) are enriched by flow cytometry and subjected to CUT&RUN with coilin, NPAT (the marker for Histone Locus Body, HLB), and IgG. (H) CUT&RUN track of the Histone Gene Clusters, with the genomic coordinates indicated at the bottom. S phase signal shown in red, G1 in green.

Specifically, when compared to the local IgG background, we detected robust coilin binding peaks at genomic loci encoding major and/or minor spliceosomal RNAs (U1, U2, U4, U5, and U12), snoRNAs (U3 and U8), and to a lesser extent, U7 snRNA (Fig. 5C-D). Demonstrating strict SMN-dependency, these prominent coilin peaks were largely abolished upon SMN depletion (Fig. 5C). Functionally, among these coilin-bound RNAs, Northern blot analysis confirmed that all tested snRNAs (U1, U2, U4) exhibited a 50% reduction in the steady state level upon SMN depletion (Fig. 5F, Extended Data Fig.4A). In contrast, there was no significant coilin enrichment across the histone locus (Fig. 5D) or at the Pol III-transcribed U6 snRNA and 5S rRNA loci^58^ (Fig. 5E), where the signal remained sparse and indistinguishable from the IgG control regardless of SMN status.

The association of coilin with histone loci is cell cycle-regulated, peaking during S phase^59^. Thus, the lack of coilin footprint at these loci in our bulk CUT&RUN was potentially due to an insufficient enrichment of S-phase cells. To address this, we engineered a stable FUCCI-5 reporter cell line^60^ to differentially label cells based on cell cycle phase, allowing us to use flow cytometry to isolate homogeneous G1 and S-phase populations (Fig. 5G). We then analyzed the CUT&RUN signal of coilin and NPAT - an essential factor for histone transcription - at the histone loci. Consistent with the literature, we observed robust NPAT occupancy at histone loci in S-phase, but not G1, cells (Fig. 5H). We also detected modest coilin enrichment at a subset of NPAT sites, specifically in S-phase cells (Fig. 5H). Because NPAT serves as the scaffold for the Histone Locus Body (HLB)-another nuclear condensate-these findings validate that our CUT&RUN approach faithfully captures native condensate-chromatin interactions under non-fixed condition.

The CB has been linked to the topological organization of chromosomes^39^, and subunits of RNA polymerase machinery have been detected within CBs^61^. This prompted us to investigate whether the nucleolar invasion of CBs impacts the global chromatin landscape. Using CUT&RUN profiling, we detected a ∼2kb enrichment of H3K4me3 flanking the transcription start sites (TSS) of coilin-bound loci, which was notably absent at coilin-free sites (Extended Data Fig.5B-C). While this suggests that coilin selectively associates with active transcription, SMN depletion did not alter the H3K4me3 at these native coilin targets (Extended Data Fig.5C). Furthermore, SMN depletion did not affect the genome-wide distribution of the active H3K4me3 or the repressive H3K27me3 mark (Extended Data Fig.5D-F). Together, these data decouple coilin’s condensate state from epigenetic regulation: while native, liquid-like coilin targets active promoters, it is dispensable for maintaining the local and global chromatin landscape, which remain unperturbed despite the liquid-to-gel hardening and intranucleolar invasion of CB remnants.

Given that native CB-bound coilin occupies a mere ∼1% of snRNA loci (Fig.5A, Extended Data Fig.5C), we sought to determine the underlying basis for this selectivity by comparing the coilin-bound and coilin-free U5 snRNA variants. At the primary RNA sequence level, we identified two differentially conserved regions (dCRs) in Stem-Loop 1 and at the Sm site - regions vital for 5’ splice site alignment^62^ and snRNA stability, respectively. We found that both dCRs share 100% identity among coilin-bound U5 snRNAs, but not coilin-free U5 variants (Extended Data Fig.5G). Importantly, coilin-bound U5s, but not coilin-free U5s, accumulate to significant levels in both the steady-state total RNA fractions and the fractions immunoprecipitated using an antibody against SmB, a core Sm-binding ring protein that confers snRNA stability and functionality^63^(Extended Data Fig.5G). At the RNA secondary structure level, Minimum Free Energy (MFE) modeling^64^ suggests that coilin-bound U5 snRNAs fold identically to the experimental model extrapolated from the cryo-EM structure^65^ (Extended Data Fig.5H). Conversely, all coilin-free U5 variants assume diverse and divergent folding states at the dCRs (Extended Data Fig.5I). This supports a model in which coilin is selectively recruited to loci producing structurally intact snRNAs, acting as a molecular sensor for snRNA integrity.

### CB-nucleolar mixing impedes telomerase activity and telomeric targeting

We have so far shown that at least two of the telomerase holoenzyme components, TCAB1 and TR, mislocalize intranucleolarly upon SMN-depletion. We next sought to determine the functional impact of CB-nucleolar mixing on scaRNP function by studying telomerase’s cellular readouts. We solubilized active telomerase RNPs into an NP40 lysate (Fig.1A) and assayed the bulk telomerase activity level using a ^32^P gel-based TRAP assay^66^ (Fig.6A). Although SMN is efficiently depleted after 3 days of dox treatment (Fig. 1A), telomerase activity remains unchanged until the sixth day, when we observe the onset of a progressive decline from 100% to below 25% of activity (Fig.6A). Importantly, the timing of the decline on day 6 coincides with the onset of the coilin-dependent proliferative slow-down and the emergence of intranucleolar CB (Fig.1C, Extended Data Fig.1E). This strict temporal correlation suggests that CB-nucleolar mixing may drive the telomerase decline.

**Fig.6.**
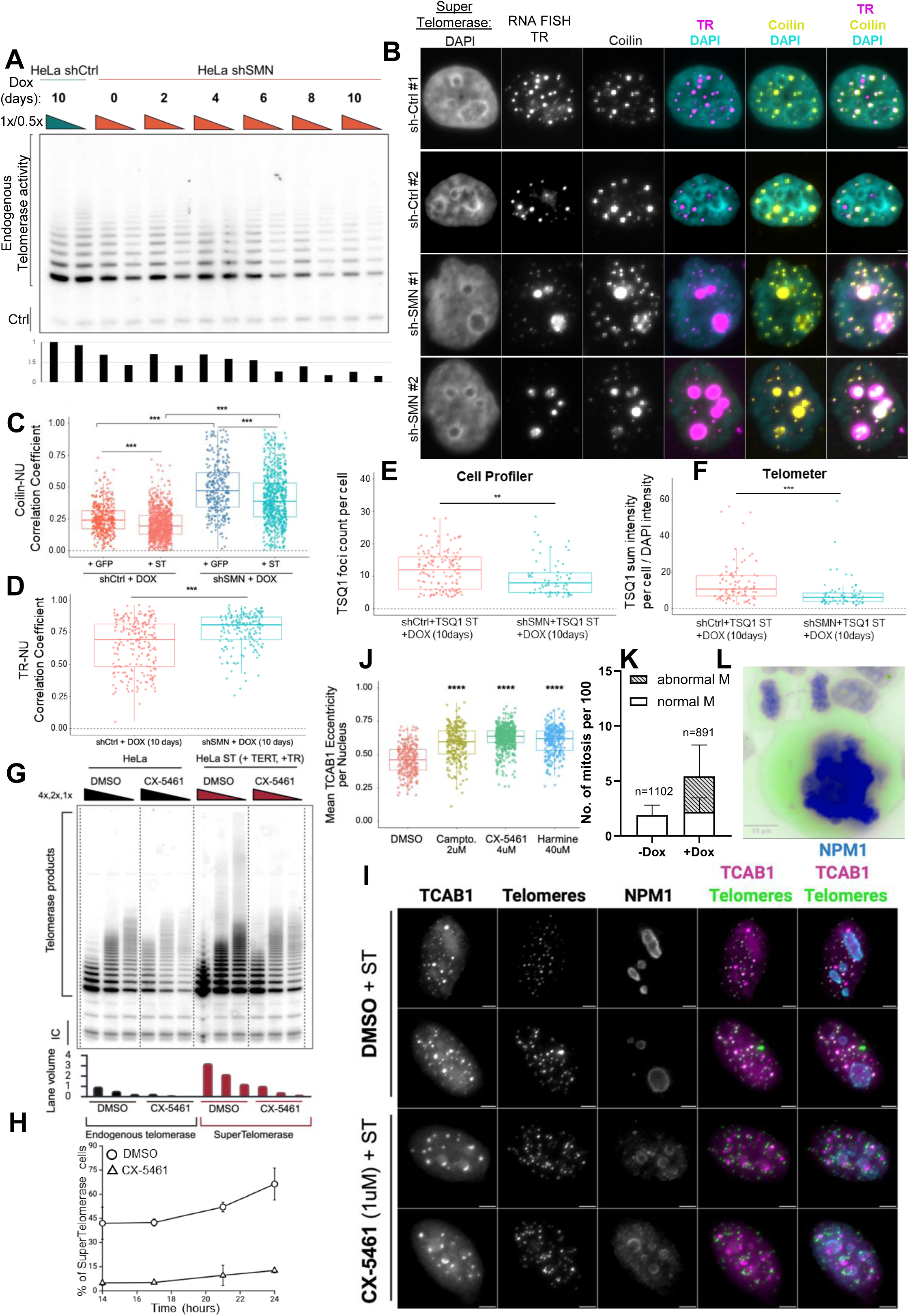
CB-nucleolar mixing impedes the targeting of coilin and telomerase scaRNP. (A) Longitudinal telomerase activity by the TRAP protocol. Active telomerase RNPs extracted/solubilized by a buffer containing 0.1% NP40 and 150mM NaCl from shSMN or shCtrl cells across the indicated time points. 1 or 0.5 µg of lysates were assayed. TRAP products quantified in the bar graph below. (B) Supertelomerase (ST) targeting to cellular telomeres: shSMN or shCtrl cells transfected with TERT and TR expressing plasmids, expressing supertelomerase. +/- DOX treatment for 10-days. The telomeric neo-CB phenotype is tracked by RNA-FISH for TR or IF for Coilin. Two representative cells are shown. Scale bar, 2 µm. (C) +/-ST condition as in B, the degree of colocalization between Coilin and Nucleolus, as marked by Fibrillarin-IF, is scored from: shCtrl + GFP (n=378), shCtrl + ST (n=915), shSMN + GFP (n=291), shSMN + ST (n=658) cells. *** p <0.001. (D) Degree of colocalization between TR and nucleolus, as marked by Fibrillarin-IF, under ST in either shCtrl or shSMN cells with 10-day dox. Data is scored from shCtrl (n=216) and shSMN (n=172) cells. ***p = 6.06e-9 (E) TSQ1 foci, reporting nascent telomere synthesis, number is quantitated by CellProfiler, in either shCtrl or shSMN cells with 10-day dox. Data is scored from shCtrl (n=92) and shSMN (n=69) cells. **p = 0.010. (F) TSQ1 foci sum intensity per cell normalized by DAPI intensity, as quantitated by Telometer is scored from shCtrl (n=92) and shSMN (n=69) cells. ***p =4.92e-7. (G) TRAP assay under the CX-5461 treatment (1uM, 24 hours), in either parental HeLa cells or HeLa ST cells. 1, 0.5, and 0.25 µg of protein lysates were assayed, with lane volume quantitated down below. (H) Longitudinal percentage of cells showing ST phenotype, as visualized in the panel I. ST-transfected cells were treated with either DMSO or CX-5461 (1uM) and imaged over time. Approximately 200 cells were quantified at each time point and condition for two biological replicates. (I) Combo telomere-FISH and co-IF characterizing the efficiency of TCAB1 targeting to telomeres in ST cells treated with DMSO or CX-5461. The nucleolar impact of CX-5461 was assessed by NPM1 staining, shown as discontinuous open rings. Successful telomeric targeting of TCAB1 shows an overlapping signal between the green and purple channels. Two representative cells are shown. Scale bar, 3 µm. (J) The mean eccentricity of TCAB1 structures was plotted to infer the TCAB1 organization upon the treatment of DMSO or indicated chemotherapy drugs at the listed concentrations. The eccentricity of TCAB1 ranges from 0 to 1, with 0 being a perfect circle. DMSO (n=282), Camptothecin (n=256, p=9.51e-27), CX-5461 (n=469, p=4.40e-70), Harmine (n=253, p=6.60e-40). (K) Quantitation of mitotic cells, either with normal or aberrant morphologies. Percentages of the two types are shown. +/- Dox treatment for 10 days. N=1102 or 891, respectively. (L) Representative examples of mitotic abnormalities in 10-day treated cells: a pair of late mitotic cells connected by a lagging chromosomes forming telomere-to-telomere bridge and an enlarged mitotic cell with polyploidy. Scale bar = 10 µm.

Having shown that intranucleolar CBs lose their genomic targeting to snRNA and snoRNA loci, we asked whether this intranucleolar entrapment also impedes their ability to target telomeres. To test this idea, we engineered a “Super-Telomerase (S-T)” system by simultaneously overexpressing TERT and TR, which together are sufficient to reconstitute active telomerase RNPs^67^. In SMN-proficient cells, these S-T RNPs heavily concentrate at the telomeres^44^. This artificially drives the entire endogenous pool of CBs to relocate to most if not all telomeres, forming quantifiable “neo-CB” foci (Extended Data Fig.6A). However, when we induced CB-nucleolar mixing by SMN depletion, the formation of these telomeric neo-CB was severely diminished. Instead, the majority of coilin and TR molecules remained spatially trapped within the nucleolus (Fig. 6B), as evidenced by their increased colocalization with fibrillarin (Fig. 6C-D).

To test whether telomere synthesis is impacted, we reconstituted a variant S-T by expressing TSQ1-TR^68^ (Tolerated SeQuence), a template-mutant TR that inserts a variant telomeric sequence, allowing the detection of *de novo* telomere synthesis. We observed a significantly reduced TSQ1 DNA-FISH signal upon shSMN induction, affecting both foci number and foci intensity (Fig. 6E-F, Extended Data Fig.6B). Thus, we demonstrated that the proper genomic targeting of CB components and the telomeric synthesis are compromised upon CB-nucleolar mixing.

As an orthogonal inducer of CB-nucleolar mixing, we treated SMN-depleted cells with the RNA Pol I inhibitor CX-5461 (Extended Data Fig.2F-J). This treatment resulted in a consistent 4-fold reduction in telomerase activity. Importantly, this reduction was observed both in the endogenous RNPs of the control cells and in cells overexpressing the S-T RNPs (Fig. 6G). Live cell imaging of endogenous Halo-TCAB1 indicated that, in untreated cells, this protein was efficiently targeted to spherical telomere foci by the S-T, with minimal contact with nucleolin-positive nucleolus (Extended Data Fig.6C). However, this S-T targeting phenotype was markedly reduced (from 44-70% to 5-10%, Fig. 6H) in cells treated with 1µM of CX-5461. Under these conditions, TCAB1 translocated to the peri-nucleolar, crescent-shaped caps that failed to overlap with telomeric DNA despite S-T overexpression (Fig. 6I), suggesting that TCAB1 targeting to telomeres was impeded by its nucleolar translocation. Furthermore, we induced a comparable degree of CB-nucleolar association by treating cells with distinct stress-inducing agents, camptothecin and harmine (Extended Data Fig.6D), evidenced by a gain of eccentricity for both TCAB1 (Fig.6J) and coilin (Extended Data Fig.6E). These results suggest that the nucleolar sequestration of telomerase components occurs broadly as a general response to nucleolar stress.

We detected an increase in the percentage of mitotic cells from 2% to 6% after 10 days of SMN depletion (Fig. 6K), suggesting an elevated level of mitotic stress. In addition, we found a marked increase in mitotic abnormalities (from 4.5% to 40.5%, Fig. 6K), including misaligned chromosomes, micronuclei, polyploidy, and notably, telomere-to-telomere end fusions (Fig. 6L and Extended Data Fig.6F). Although mitotic failure can be induced by general DNA damage during SMN loss^69^, the emergence of telomeric end fusions specifically points to a concurrent failure in telomere maintenance^70^. Thus, we propose that the nucleolar sequestration of telomerase, resulting from CB-nucleolar mixing, precipitates telomere-driven genomic instability.

## DISCUSSION

Here, we propose an inter-condensate crosstalk mechanism by which spliceosome stress is communicated intranucleolarly via CB-nucleolar intermixing, where remnants of the dissolved CB undergo a liquid-to-gel hardening process while gaining occupancy at the rDNA chromatin. The resulting intranucleolar Coilin directly suppresses ribosomal biogenesis *in cis* and telomerase functions at telomeres *in trans*.

Our findings demonstrate direct and functional crosstalk between two distinct phase-separated condensates, the CB and nucleolus. Coilin relocation to the nucleolus has been observed as a stress marker and attributed to passive sequestration. Our results suggest an active role in which intranucleolar coilin gains stable, specific, and functional interactions at the peri-FC zone surrounding rDNA chromatin, culminating in reduced nascent rRNA production. In this new model, coilin plays a tripartite role as a spliceosome-integrity sensor, an inter-condensate messenger, and an rDNA transcription dampener.

We found that the intranucleolar phase transition of non-nucleolar proteins directly and simultaneously impacts multiple RNP functions. The material properties of the nucleolar core (FC) resemble those of a viscoelastic gel with restricted mobility^6^. When CB proteins are mistargeted to either the peri-FC upon SMN depletion or the nucleolar rDNA caps upon CX-5461 treatment, their liquid-to-gel hardening enhances coilin’s dwelling time on rDNA, thereby converting coilin into a stable rDNA repressor impeding transcription^41^. In addition, hardening of TCAB1 - the telomerase activity switch^43,44^ - would limit the productive assembly between TERT and TR. Furthermore, intranucleolar mislocalization of TCAB1 can cripple telomerase’s telomeric targeting mechanism^13^ by intranucleolar sequestration of telomerase^28^, leading to compromised telomere synthesis and chromosomal integrity.

Our data support a previously unappreciated role for SMN as a general solubility factor for CB components, including the active telomerase RNP, licensing their extrication from the entangled chromatin matrix. This raises the question of how SMN promotes the low-viscosity, liquid-like state of CBs while simultaneously preventing intermixing with the dense nucleolus. While both CB and nucleolar-residing RBPs tend to harbor RG/RGG boxes at high frequency^49^, only CB residents possess the sDMA-modified RG box that is selectively recognized by the SMN Tudor domain^71^. Indeed, Coilin mislocalizization to the NU occurs in parallel with its simultaneous loss of sDMA modification^25^. Thus, we propose that the engagement between SMN and sDMA physically masks arginine residues, preventing them from promiscuously binding to RNA via base stacking and hydrogen bonding^72^. By shielding these arginine motifs, SMN maintains the liquid-like mobility and solubility of CB RBPs^73^. Without this protection, these unmasked arginines would drive their entropic entanglement into RNA-dense condensates^74^, such as the nucleolus.

Our evidence further supports a new role for coilin as a chromatin-associated RNP suppressor. We demonstrate that coilin depletion sufficiently restores molecular and cellular defects induced by SMN depletion, supporting a model wherein coilin antagonizes SMN’s general solubility role by driving RNP hardening, consistent with the finding that CBs are transcriptionally inert^75^. Mechanistically, coilin possesses a general, sequence-independent affinity for RNA^37^ and a high capacity for self-oligomerization^35^, rendering it an aggregation-prone RNA nucleator – particularly when unchaperoned by SMN. Furthermore, our CUT&RUN data captures coilin’s footprints at the 3’ readthrough region of the transcription units encoding snRNA, snoRNA, and rRNA. These regions are typically co-occupied by stalled RNA Polymerase I or II^40,76^ and R-loops^77^ – structures that require fluid dynamics for efficient resolution^78^. It is tempting to speculate that the CB may normally function as a “transcription termination condensate” that facilitates the release of nascent RNAs^76^. In SMA, we speculate that the aberrant hardening of coilin at the rRNA 3’ site freezes the termination machinery, effectively locking the pre-rRNAs on the chromatin template and slowing down their further processing.

Finally, our findings position SMA within the emerging class of “condensatopathies”, identifying it as a prototype of “inter-condensate mismanagement” – a novel pathogenic mechanism driven not by the failure of a single condensate, but by the breakdown of immiscibility between two distinct nuclear organelles. Moreover, many causal mutations have been mapped to the Intrinsically Disordered Regions (IDRs) of nuclear condensate-associated genes, such as TCOF1 in Treacher Collins syndrome^79^, NPM1 in leukemia^80^, TCAB1 in dyskeratosis congenita^42^, and TPP1 in aplastic anemia^81^. Our work also provides a mechanistic framework for understanding how such inter-condensate mismanagement can drive the etiology of both Ribosomopathies and Telomereopathies. Furthermore, the ability of the Cajal body and nucleolus to maintain their spatial segregation serves as a sensitive biomarker for nuclear homeostasis.

Our model suggests that therapeutic strategies need not strictly target the missing condensate-associated gene but could instead aim to enforce the biophysical boundary between disrupted condensates. Small molecules that modulate the viscoelasticity of the nucleolus or enhance the liquid-like mobility of CB remnants could potentially reverse the pleiotropic defects of SMA^82^-restoring ribosome biogenesis and chromosomal integrity in one stroke.

## Acknowledgements

The work is indebted to Grazia Raffa (Sapienza University of Rome) for helpful discussion; Luke D Lavis (Janelia, HHMI) for generous support with Halo ligands and chemistry; and Jens Schmidt (Michigan State) for sharing the Halo-TCAB1 HeLa cell line and advice. We would like to thank the core facility support from Sharon Connelly, Michelle Townsend, and Kerry Campbell (Cell Culture), Andrey Efimov and Neil Johnson (Biological Imaging), Jodina Hazel (Glass Washing/Medium Prep), Manna Ahmed and Johnathan Whetstine (Genomics Resource), James Oesterling and Joan Font-Burgada (Cell Sorting), Cynthia Meyers (Organic Synthesis), Jessica Rodgers, James Hambor, and Kendall Berry (Radiation Safety) at FCCC, and Anneliese Faustino and Hsin Yao Tang (Proteomics and Metabolomics) at Wistar Institute. We also thank Diane Jacquindo, Ariana Vargas, Christina George, John Gricoski, and David Wiest for administrative support, as well as Amanda Purdy, Sarah Whealen, and Glenn Rall for support in academic affairs, Xavier Graña and Eishi Noguchi for graduate program support, and Siddharth Balachandran for feedback. Grant support for this study includes NIH/NIGMS grants GM150538 and GM150538-02S (NanoLive) to L.C., subcontracts HL172961, AI164333, AG065204, CA269660 to L.C., NSF 2334246 and NSF 2334624 to G.L., R01NS102451 to L.P., and NIH/NCI Cancer Center Support Grant P30 CA006927. J.H. is supported by NIH T32 GM142606.

## Author Contributions

L.C. conceived the study, prepared the figures, and wrote the manuscript with input from the authors. L.C., A.D.M., and J.H. conceptualized and designed the study. A.D.M. and J.H conducted, analyzed, and interpreted the majority of the experiments, with A.D.M. responsible for all genomic, bioinformatics, and most of the nascent rRNA efforts, J.H. for telomerase/telomere characterizations and all fix-cell imaging, and V.N. for live confocal and nanolive imaging, and the generation of the necessary stable cell lines. C.G.L. contributed to the initial phenotypical discovery; E.P., to the proteomic and the nascent RNA pulse-chase experiments; M.R., to cell cycle analyses; B.S., to northern blotting and reagents; A.E., V.N., and L.C., to FRAP; Y.T and G.L, to 3D construction and quantification of confocal images; C.M. and J.H., to the synthesis of Halo ligands; E.G. and L.P, to reading, conception, and feedback; L.S. and L.P., for the inducible SMN RNAi cell line construction.

## METHODS

### Inducible HeLa shSMN Cell Culture

HeLa cells were cultured in Dulbecco’s modified Eagle’s medium (DMEM) with high glucose (Invitrogen) containing 10% fetal bovine serum (HyClone), 2mM L-glutamine (GIBCO), and 1X penicillin/streptomycin (GIBCO). RNAi was induced by addition to the growth medium of Doxycycline HCl (Fisher Scientific) at the final concentration of 100 ng/ml. Antibiotic selection was carried out with 5 μg/ml Blasticidin-S hydrochloride (Invitrogen), and 5 μg/ml Puromycin (Sigma).

### Lentiviral Constructs and Viral Production for shSMN cell line

The pLenti6/TR (Invitrogen) vector constitutively expresses the tetracycline-dependent repressor (TR) protein under the control of the CMV promoter as well as the blasticidin resistance gene from the SV40 promoter. The pLenti.pur/SMN_RNAi_ construct expresses an shRNA targeting human SMN mRNA (5’-GAAGAAUACUGCAGCUUCC-3’) under the control of a tetracycline-regulated H1TO promoter as well as the puromycin resistance gene from the PGK promoter. This vector was generated by cloning annealed oligonucleotides corresponding to the SMN_RNAi_ hairpin sequence into pSUPERIOR.puro (Oligoengine) followed by transfer of the fragment containing the H1::SMN_RNAi_ and PGK::puromycin cassettes into the pRRLSIN.cPPT.PGK-GFP.WPRE vector (Addgene plasmid 12252) as a lentiviral backbone. Viral stocks pseudotyped with the vesicular stomatitis G protein (VSV-G) were prepared by transient co-transfection of 293T cells using the ViraPower Lentiviral Packaging Mix (Invitrogen) following manufacturer’s instructions.

### Inducible HeLa shSMN generation

The HeLa-SMN_RNAi_ cell line was generated through two consecutive rounds of lentiviral transduction. First, HeLa cells were transduced with the plenti6/TR vector and stable cell lines established by antibiotic selection with 5 μg/ml Blasticidin-S hydrochloride (Invitrogen) and cloning by limiting dilution. A resulting clone with high expression of TR was subsequently transduced with the pLenti.pur/SMN_RNAi_ vector followed by antibiotic selection with 5 μg/ml Blasticidin-S hydrochloride (Invitrogen) and 5 μg/ml Puromycin (Sigma) and cloning by limiting dilution. The HeLa-SMN_RNAi_ cell line was selected among various stable clones after screening for normal SMN expression in the uninduced state and the greatest level of regulated SMN knockdown upon doxycycline addition.

### Protein Extraction

Protein extraction was performed using a sequential lysis approach to separate soluble and insoluble protein fractions. HeLa shSMN cells subjected to 9-12 days of DMSO (-Dox) or doxycycline (+Dox) were harvested in ice-cold PBS, pelleted at 500 × g for 5 minutes, and resuspended in ice-cold NP-40 lysis buffer (10 mM Tris-HCl pH 7.5, 150 mM NaCl, 1% NP-40, 1× protease inhibitor cocktail). Cells were incubated at 4°C for 15 minutes under rotation and lysates were cleared by centrifugation at 21,000 × g for 5 minutes at 4°C. The resulting supernatant was collected as the soluble protein fraction.

The remaining insoluble pellet was resuspended in RIPA buffer supplemented with a 1:500 dilution of DNase I (50 mM Tris-HCl pH 7.5, 150 mM NaCl, 1% NP-40, 0.5% sodium deoxycholate, 0.1% SDS, 1 mM EDTA, 1× protease inhibitor cocktail) to solubilize chromatin-bound and nuclear matrix-associated proteins. After a 30-minute incubation at room temperature under rotation, samples were centrifuged again at 21,000 × g for 15 minutes, and the supernatant was collected as the insoluble protein fraction.

### Western Blot

Equal amounts of protein, as quantified by BCA Protein Assay (Thermo Fisher Scientific), were mixed with SDS sample buffer (Invitrogen) and Reducing Agent (NuPage) and incubated at 70°C for 10 minutes. Proteins were separated on 4–12% Bis-Tris SDS-PAGE gels (Invitrogen) and transferred to 0.45 µm PVDF membrane (Millipore) for 1 hour at 100V using a wet transfer system. Membranes were blocked in TBS blocking buffer (Licor) at room temperature for 1 hour, followed by incubation with primary antibodies overnight at 4°C (table?). After washing, membranes were incubated with IRDye 680RD-conjugated anti-mouse and IRDye 800CW-conjugated anti-rabbit secondary antibodies (Licor) for 1 hour at room temperature. Visualization was carried out by detecting the secondary antibody’s fluorescent signal using the Odyssey M Imager (Licor). Band intensities were quantified using Image Studio and normalized to the total protein.

### Mass Spectrometry

For LC-MS/MS analysis, proteins were prepared as described under the western blot methods, run for approximately 1/2 centimeter into an 8% Bolt gel (Invitrogen) and stained with Imperial Protein Stain (ThermoScientific, Cat. No. 24615) following the manufacturer’s instructions. The gel was provided to our collaborators at the Wistar Institute’s Proteomics and Metabolomics Facility, where they took the gel and subjected it to reduction (TCEP) and alkylation (IAA) of cystines followed by trypsin in-gel microdigestion.

The digest solution was analyzed using a 40-min run on a nano LC coupled to an Orbitrap Astral mass spectrometer (Thermo) in Data-Independent Acquisition mode. Peptide identification was performed by searching against a spectral library generated from the SwissProt Homo sapiens and a contaminant database using DIA-NN v1.9.2. The “match between runs” feature was enabled, and the precursor, peptide and protein false discovery rates (FDR) were set at 1%. Abundance values were normalized using the MaxLFQ algorithm, fold changes were calculated using the normalized abundance values and missing values were replaced with the minimum value of the dataset, when applicable.

### Immunofluorescence Staining and Fluorescence In Situ Hydridization (FISH)

Cells were seeded into a 24-well plate containing sterile glass coverslips (Fisher, 12-541-002) and cultured for 1-2 additional days before fixation. Cells were then permeabilized with 500uL sucrose-triton buffer (20mM HEPES ph = 7.9, 50mM NaCl, 300 mM sucrose, 3mM MgCl2, 0.5% Triton-X100) for 2 minutes at room temperature, rinsed twice with 500uL PBS (Ca2+/Mg2+ free), and fixed in 500uL 10% formalin (for 10 minutes at room temperature. Two additional PBS washes were performed and immediately followed by the staining protocol or optional storage (4°C in 70% ethanol). Cells were re-permeabilized with 500uL 0.1% Triton X-100 in PBS for 15 minutes, nutating at room temperature, followed by two PBS washes. The coverslips were then blocked with 250uL 1% BSA in PBS for 30 minutes, nutating at room temperature. Primary antibodies were diluted 1:1000 in blocking solution, and 250uL were added per well for an overnight incubation at 4°C. Following primary antibody incubation, the coverslips were washed 3x for 5 minutes each wash with 500uL PBS. Coverslips were incubated with 250uL Alexa Fluor–conjugated secondary antibodies (Thermo Fisher Scientific), diluted 1:10,000 in blocking solution for 45 minutes, nutating at room temperature. This was followed by three PBS washes. If not proceeding to FISH, the coverslips were mounted using ProLong Gold Antifade Mountant DAPI (Invitrogen P36935). If proceeding to FISH, the secondary antibody signal was fixed with 500uL of 10% formalin, pre-warmed, nutating for 10 minutes. The coverslips were progressively dehydrated with ethanol incubations: 70% 95% 100%, 5 minutes each, stationary. After the last incubation, the coverslips were airdried for 10 minutes. For DNA FISH, the coverslips were inverted onto 20uL droplets of 0.1uM probe (Cy5-TelC or Cy3-TSQ1), diluted in DNA FISH hybridization buffer (10mM Tris-HCl pH = 7.4, 70% formamide, 0.5% Roche blocking reagent). They were then denatured on a heating block at 80C for 5 minutes before being incubated in a dark humidity chamber overnight at 4C. The coverslips were then washed 2x, 15 minutes, nutating at room temperature with FISH wash buffer (2x SSC, 50% formamide), and then mounted on slides. For RNA FISH, the coverslips were rehydrated following airdrying with 500uL FISH wash buffer for 5 minutes. They were then pre-hybridized for 1 hour at 37C, nutating with 250uL RNA FISH hybridization buffer (2x SSC, 50% formamide, 100mg/mL dextran sulfate, 1mM VRC, 0.5 mg/mL salmon sperm DNA, 1 mg/mL BSA, 0.125 mg/mL yeast tRNA). RNA FISH probes (Cy5-TR) were equally mixed and diluted to 1.5ng/uL in RNA FISH hybridization buffer, and the coverslips were inverted onto 20uL of the diluted probe. They were then incubated overnight in a tissue culture incubator at 37C. The following day, the coverslips were washed 2x, 30 minutes, nutating at 37C with FISH wash buffer, before being mounted.

### Generation of stable HeLa cell lines expressing fluorescently tagged Nucleolar and Cajal Body proteins

HeLa S3 cells under shSMN (or sh control, where indicated) background at ∼1×10 confluency were used to generate stable cell lines expressing eGFP-tagged nucleolar and Cajal body proteins. For eGFP-Fibrillarin (Addgene 26673, a gift from Sui Huang), eGFP-Nucleolin (Addgene 28176, a gift from Michael Kastan), and eGFP-Coilin (Addgene 369404, a gift from Greg Matera) lines, cells were transfected with 2 µg of the respective peGFP plasmids, and after 48 h, selected with 500 µg/mL Neomycin for 10 days. GFP-positive single cells were sorted by FACS into 96-well plates containing DMEM (10% FBS) with 80 µg/mL Neomycin, and clones were validated by western blotting and fluorescence microscopy. For dual-labeled lines (eGFP-Fibrillarin/Coilin-mCherry and eGFP-Nucleolin/Coilin-mCherry), Coilin-mCherry was integrated via PiggyBac transposition by co-transfecting 1.5 µg piggybac Coilin-mCherry and 1.5 µg transposase plasmids using Lipofectamine 2000, followed by selection after 48 h with 500 µg/mL Hygromycin and 80 µg/mL Neomycin for 4 days. Double-positive cells were sorted into 96-well plates containing DMEM (10% FBS) with 80 µg/mL Hygromycin and 80 µg/mL Neomycin, and clones were validated by western blotting and fluorescence microscopy. For Halo-TCAB1 tagging in the eGFP-Coilin background, CRISPR/Cas9-mediated genome editing was performed by transfecting 1 µg sgRNA/Cas9 (Addgene #207611) and 1 µg HDR donor plasmid (Addgene #207535) using Lipofectamine 2000, followed by selection with 1 µg/mL Puromycin for 5 days. Correct insertion was confirmed by PCR using primers flanking the 5′ homology arm and Halo tag region. To excise the puromycin cassette, cells were transfected with 1 µg Cre-mCherry plasmid (Addgene #212105), and after 48 h, mCherry-positive single cells were sorted into 96-well plates by FACS. Clones were validated for Halo-TCAB1 expression by JF646 Halo ligand labeling, fluorescence microscopy, and western blotting.

### Live-cell Fluorescence recovery after photobleaching (FRAP)

Cell growth conditions: HeLa S3 cell lines stably expressing eGFP coilin under sh and shSMN backgrounds or HeLa S3 - eGFP Fibrillarin/Coilin mCherry under SMN background or HeLa S3 - eGFP Nucleolin/Coilin mCherry under shSMN background were grown in 10 cm dishes containing DMEM complete media and 80 µg/mL Neomycin or 80 µg/mL neomycin and 80 µg/mL Hygromycin. The cell lines were either treated or untreated with 1X Doxycycline to induce the expression of short hairpin (sh) for 9 days with media and doxycycline replaced every 2 days. On day 8, cells were seeded in a µ-slide 8-well cell culture chamber in DMEM complete media containing 80 µg/mL Neomycin or 80 µg/mL Neomycin and 80 µg/mL Hygromycin.

Cell growth conditions for HeLa S3 – eGFP Coilin/Halo TCAB1: HeLa S3 – eGFP Coilin/Halo TCAB1 under shSMN background was grown in 10 cm dish containing DMEM complete media and 80 µg/mL Neomycin. The cells were either treated or untreated with 1X Doxycycline to induce the expression of short hairpin (sh) for 9 days with media and doxycycline replaced every 2 days. On day 8, cells were seeded in µ-slide 8-well cell culture chamber in DMEM complete media containing 80 µg/mL Neomycin and 200nM janelia flour 646 (JF 646).

Fluorescence Recovery After Photobleaching (FRAP) was carried out using Leica TCS SP8 FSU laser scanning confocal microscope with 63x oil lens. Briefly, the entire normal Cajal body (CB) or the entire aberrant CB (cap-like, ring-like, or patchy) was bleached with 100% 488 nm laser for 10 frames, and the fluorescence recovery frames were collected every 20 seconds for 480 seconds in a z-stack with excitation and emission maxima at 488 nm and 513 nm, respectively. The Z-stack images were collected using LAS X (Leica Microsystems) software.

Quantification: First, fluorescence intensities in Z-stack images were summed using Zprojection in ImageJ. Then the fluorescence intensities of bleached and non-bleached CBs of time lapse images were quantified using CellProfiler. Non-bleached CB served as a control and was always selected from the same cell as bleached CB. Normalization of the quantified data was done in three steps i) percentage of control CB (non-bleached) bleaching was calculated by dividing the fluorescence intensity of control CB at each time frame by the fluorescence intensity of control CB in the pre-bleached time frame. ii) fluorescence intensities of bleached CBs were normalized by dividing fluorescence intensity of bleached CBs at each time frame by the percentage of control CB bleaching at the same time frame. iii) Double normalization of bleached CBs was performed by using the below formula

□□□□(□)=Ref_pre−bleach_/ref(t)*FRAP(t)/FRAP_pre−bleach_

Ref_pre−bleach_ = mean intensity of reference region pre bleach

Ref(t) = Intensity of reference region at time point t

FRAP(t) = intensity of FRAP region at time point t

FRAP_pre−bleach_ = mean intensity of FRAP region pre bleach

### 4SU pulse chase

To label newly synthesized RNA, cells were incubated with 4-thiouridine (4sU; Sigma-Aldrich, Cat. No. T4509) at a final concentration of 500 µM or DMSO (no pulse control) for 1hr at 37°C in complete growth medium. After labeling, cells were washed with PBS and immediately processed for total RNA extraction, with the exception of the 8hr time point, which was subjected to complete growth medium with 2.5 mM uridine and left to incubate at 37°C for 8 hours after the initial 1hr 4SU pulse. Labeling was performed at time points of 0 hrs and 8 hrs, as described, to assess RNA transcription rates and turnover dynamics.

### Nascent RNA isolation

For enrichment of 4sU-labeled (nascent) RNA, biotinylation of 4sU-incorporated RNA was performed by incubating total RNA of the same concentration with 0.5 mg/mL EZ-Link HPDP-Biotin (Thermo Fisher, Cat. No. 21341) in 10X biotinylation buffer (100 mM Tris pH 7.4, 10 mM EDTA) for 1.5 hours at room temperature. Biotinylated RNA was phase-separated twice using chloroform, precipitated using isopropanol and NaCl, washed with 75% ethanol and resuspended in RNase-free water. The extracted RNA was purified using streptavidin-coated magnetic beads (Miltenyl Biotec) and after heating the total RNA for 10 minutes at 65°C and incubating the denatured RNA with the beads for 15 minutes before introducing them to the column (μMacs, Miltenyl Biotec), the bound RNA was subjected to three washes of wash buffer at 65°C (100 mM Tris pH 7.4, 10mM EDTA, 1M NaCl, 0.1% Tween 20) followed by three washes of room temperature wash buffer and the elution of newly transcribed RNA with two rounds of 100 mM DTT into buffer RLT (RNeasy MinElute Cleanup Kit, Qiagen). Extraction was followed by RNA cleanup using the aforementioned kit, following the manufacturer’s protocol.

The concentration, quality and integrity of both total and labeled RNA fractions were assessed using Qubit analysis and TapeStation (Agilent Technologies). RNA was stored at −80°C until further use.

### Biotinylated RNA Dot Blot

Biotinylated RNA was measured using a NanoDrop One microvolume UV-Vis spectrophotometer, normalized to the same concentration and serially diluted by 10-fold. After activating the positively charged nylon membrane (Hybond-N+, Amersham) using nuclease-free water and air-drying for 5 minutes, the samples were spotted directly onto the membrane alongside a 5’ dual biotinylated oligo standard (pRMS217, IDT, 5’52-Bio/GAC GCT GCC GAA TTC TAC CAG T), using manual pipetting. Membranes were air-dried for 5 minutes before UV-crosslinking the RNA to the membrane at 0.2 J/cm2 (254nm). After immobilizing the RNA, the membrane was blocked in a blocking solution [10%SDS, 1mM EDTA in PBS] for 20 minutes at room temperature.

For nascent RNA detection, membranes were incubated for 15 minutes with a 1:50,000 dilution of 1mg/mL streptavidin-horseradish peroxidase (HRP) (Thermofisher) in blocking solution, protected from light. While continuing to protect the membrane from light, two washes were performed using the blocking buffer solution, followed by two washes of wash buffer I (1% SDS, 1mM EDTA in PBS) and two washes of wash buffer II (0.1% SDS, 1 mM EDTA in PBS) for ten minutes each. The biotin-bound HRP’s chemiluminescence was developed using an enhanced chemiluminescence (ECL) western blotting substrate (Pierce) and visualized using the Licor Odyssey M. Signal intensities were quantified using Image Studio.

### TapeStation

After nascent RNA isolation, the recently transcribed RNA was assessed for its integrity and concentration using the Agilent Model 4150 TapeStation system (Agilent Technologies, Santa Clara, CA, USA) with the High Sensitivity RNA ScreenTape assay, following the manufacturer’s instructions. Briefly, 2 µL of each RNA sample, mixed well with 1uL of sample buffer, was loaded onto the RNA ScreenTape alongside a ladder. Electrophoretic separation and fluorescence detection were carried out automatically by the TapeStation system.

RNA integrity was evaluated using the RNA Integrity Number equivalent (RINe), with values ≥7.0 considered suitable for downstream applications such as reverse transcription and qPCR. RNA concentrations were determined based on the electropherogram peak areas and confirmed to fall within the quantifiable range of the assay. All samples were run in a single batch to minimize technical variation.

### CB solubility Quantitation by LC/MS

To characterize the biochemical solubility of Cajal body components upon CB-NU mixing, we solubilized NP40-insoluble chromatin pellets with a high-salt DNase I-containing RIPA buffer and analyzed the relative protein abundance by LC-MS/MS. Most CB proteins, according to a Coilin interactors survey^22^, showed significantly reduced abundance in the Coilin-enriched chromatin fraction (n=53, p-values adjusted to account for multiple testing using Benjamini-Hochberg FDR correction, Fig.2E), with only a few significant exceptions (n=4). This reduction occurred broadly across diverse nuclear RNA pathways involved in snRNA and rRNA biogenesis, mRNA transcription and processing, cell cycle progression, and DNA repair and replication (Fig. 2E), including Coilin, Little Elongation Complex (LEC)^40^, RNA processing enzymes TOE1, TGS1, and H/ACA scaRNP components.

### Nano-live microscopy

HeLa S3 cells stably expressing eGFP-Coilin under an shSMN background were cultured in DMEM supplemented with 10% fetal bovine serum (FBS) and 80 µg/mL neomycin. To induce SMN knockdown, cells were treated with doxycycline (1X) for 4 days in 10-cm dishes. At the end of day 4, cells were reseeded onto 35-mm µ-dishes (ibidi #80136) at ∼40% confluency in DMEM containing 10% FBS, 80 µg/mL neomycin, and 1× doxycycline.

On day 5, live-cell imaging was performed using a NanoLive holotomographic microscope under controlled conditions (37 °C, 5% CO^2^). Time-lapse imaging was carried out over an extended period, with brightfield images acquired every 60 min and GFP fluorescence images acquired every 120 min. Acquired 3D image datasets were analyzed using NanoLive Browser software, and image stacks containing optimal brightfield and fluorescence information were selected, stacked, and exported as TIFF files. Subsequent image processing and analysis were performed using ImageJ software.

Control cells not treated with doxycycline were imaged in parallel under identical conditions.

### Telomerase Repeat Amplification Protocol (TRAP)

To measure telomerase activity, a two-step TRAP procedure was performed according to (Kim and Wu, 1997). NP-40 cell lysates were measured by protein Bradford assay and normalized across experimental samples. Sequential dilutions of 1ug, 0.5, and 0.25ug of lysate were incubated with telomeric primers for a 30 min initial extension step at 30 C in a PCR machine, followed by 5 min of inactivation at 72C. 0.5uL of product from the extension step was then used in a PCR reaction (24 cycle of 30s at 94C, followed by 30s at 59C) in the presence of 32P end-labeled telomeric primers before 10min at 72C. 32P telomere primers were labeled using T4 PNK protocol and purified by a micro-spin G-25 column (cytiva, 27-5325-01). The TRAP PCR reactions were resolved by 8% polyacrylamide gel electrophoresis running at 500V for 45 minutes, room temperature. The gel was exposed to a phosphor-imaging screen overnight and scanned by a Typhoon scanner. The lane volumes of scanned image were quantitated using the ImageQuant TL software (cytiva).

### SuperTelomerase Assay

Cells were transfected with a 3:1 ratio of pBS-U1-TR to pcDNA-3F-TERT following standard Lipofectamine 2000 procedures. For detection of newly synthesized telomeres by DNA FISH, pBS-U1-TSQ1 was used in replacement of the WT TR plasmid. shSMN cells were transfected on Day 7 of DOX treatment and fixed on Day 10, before undergoing staining procedures: IF, RNA FISH, DNA FISH (as described above). CX-5461 treated cells were given 1uM of compound 4 hours prior to ST transfection and were fixed 48 hours post-transfection. The live-cell SuperTelomerase assay was performed by seeding 10,000 HeLa-HaloTCAB1 cells in an Ibidi 8 chamber glass-bottom coverslip (#80806). Cells were seeded in DMEM containing 200nM of JF552. The next day, DMSO or CX-5461 (1uM) was added to the chambers. 4 hours after compound addition, the cells were transfected with SuperTelomerase and GFP-nucleolin. Images of HaloTCAB1 (approx. 200 cells each time-point) were taken at hours 14, 17, 21, and 24 post-transfection. Cells with small, dispersed round TCAB1 foci (indicative of telomere targeting) were counted as being + ST cells, while those with large, residual CB-like foci or drug-induced crescent-shaped nucleolar capping structures were counted as non-ST.

### 3D model construction and quantitation

Four channels were extracted from the raw LIF file: blue (DAPI), yellow (FC, RPA194), cyan (DFC, fibrillarin), and magenta (CB, coilin). To account for anisotropic voxel spacing in the microscopy data, anisotropic Gaussian smoothing was first applied, with weaker smoothing in the lateral directions and stronger smoothing along the axial direction to reduce z-axis noise while preserving in-plane structural details. The yellow, cyan, and magenta volumes were then segmented using watershed with H-maxima suppression to separate touching cells and obtain well-defined 3D object masks. Based on these segmented volumetric regions, 3D surface meshes were generated using the marching cubes algorithm, which extracts isosurfaces from voxelized data as triangulated geometric representations. This pipeline converts multichannel fluorescence image stacks into instance-resolved 3D surface models for visualization and quantitative analysis.

The violin plot in Fig.3E shows the distribution of yellow objects that overlap with either the cyan or magenta channels. To quantify these overlaps, connected objects in each channel were first identified after a dilation step. Dilation was applied to the yellow channel to bridge small gaps and merge closely adjacent regions, reducing the chance that a single object would be fragmented and counted as multiple objects because of segmentation imperfections or image noise. Connected-component labeling was then used to assign a unique ID to each object. The same procedure was applied to the cyan and magenta channels to identify the corresponding sets of labeled objects. Overlap was determined by comparing pixel coordinates among the labeled regions, and the resulting counts of overlapping yellow objects were summarized in the violin plot.

The histogram (Extended Data Fig.3B) summarizes, for each slice in the z-stack, the percentage of yellow-channel pixels that overlap with the magenta channel and the percentage that overlap with the cyan channel. Overlap was calculated by first counting the number of yellow-channel pixels co-localized with the magenta or cyan channels, and then normalizing by the total number of yellow-channel pixels in that slice. To further quantify the fraction of the yellow channel overlapping with either magenta or cyan, the union of the magenta and cyan overlapping pixels was divided by the total number of yellow-channel pixels. These normalized values were then plotted as a histogram.

### CUT & RUN

Cleavage under targets release under nuclear (CUT and RUN) was performed using Epicypher CUTANA™ ChIC/CUT&RUN Kit version 4.0 (SKU 14-1048). In accordance with kit instructions, when possible, for each cellular condition 250,000 HeLa cells per antibody were harvested and washed twice to remove debris. Then cell conditions were split into .2mL tubes 250,000 cells per antibody. Magnetic beads coated with concanavilin A were activated by two washes in activation buffer and allowed to bind to cells for 10 minutes at room temperature. The cells were then switched to antibody buffer which contains 2mM EDTA and .01% digitonin and incubated overnight with primary antibody. The next day cells were washed twice with cell permeabilization buffer and incubated with pAG-Mnase for 10 minutes room temperature to allow binding. Calcium was then added to 2mM and reactions were incubated at 4C on a nutator for 2 hours. Reactions were stopped by the addition of STOP buffer and heated at 37C for 10 minutes. Cells were allowed to bind to magnetic stand and the supernatants were removed and purified using Ampure XP beads at a 1.8X bead ratio. Cleaned supernatants were stored at –20C before library prep. Library prep was done using the NEB Ultra II kit (NEB E7645L) with single index barcodes (NEB E6609S). PCR amplification was done according to Epicypher’s recommendation combining annealing and extension steps to selectively amplify shorter fragments. Sequencing was done through Azenta pair ended 150bp to an average depth of 20 million reads per antibody.

### CUT & RUN Analysis

Reads were demultiplexed by Azenta. We performed analysis both on the Fox Chase Cancer Center (FCCC) computational cluster Zorro, and on the methylome.ai cluster of Dr. Hayan Lee at FCCC. Reads were aligned to a custom version of the Hg38 genome which contains an annotated version of an rDNA repeat (Subin et al. 2023, Journal of Biological Chemistry) using Bowtie2. RPGC bigwigs and heatmap plots were generated using Deeptools. Peaks were called using MACS3 and analyzed in R using DiffBind. Code is available upon request.

---

**Fig.S1.**
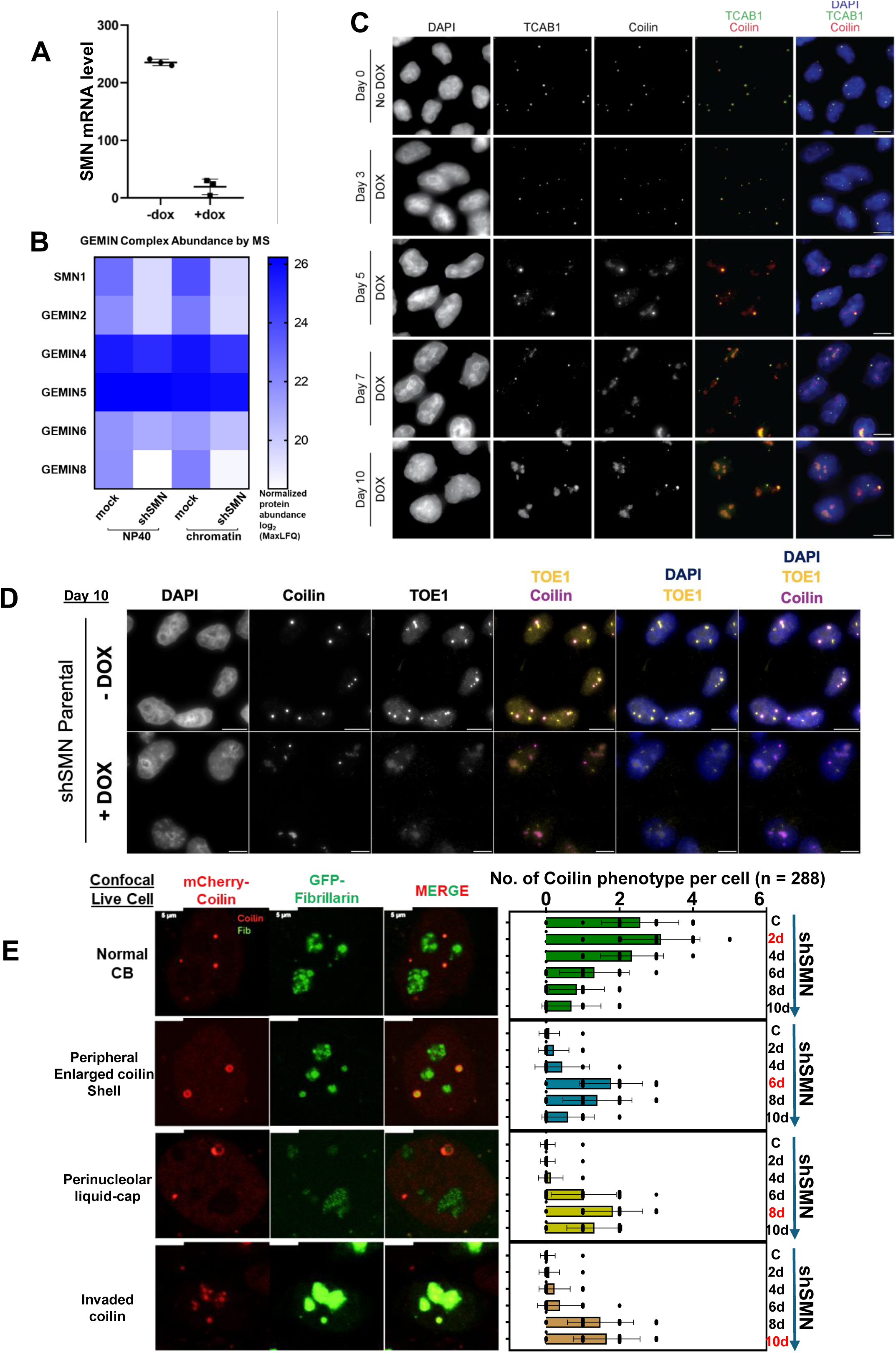
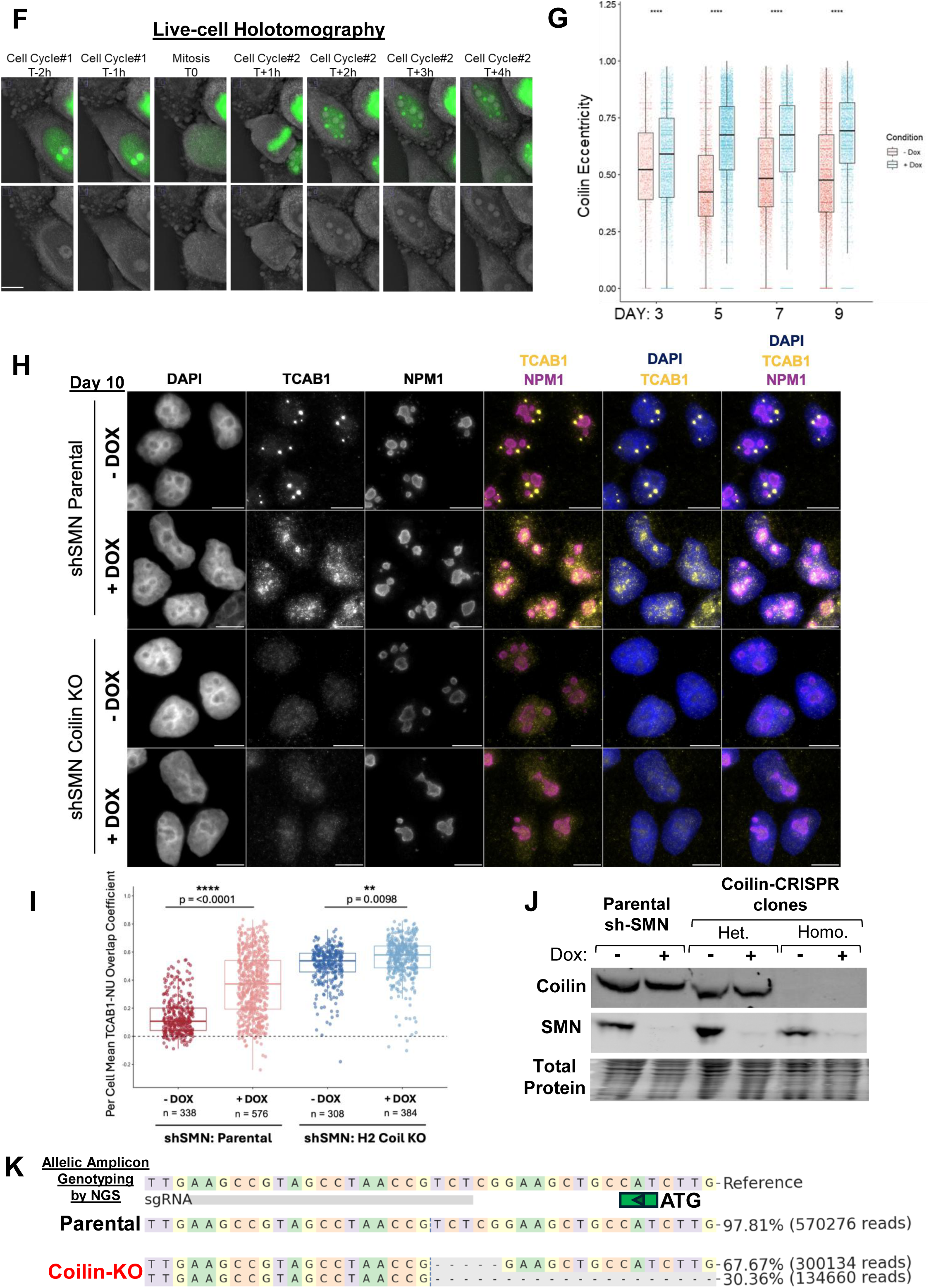
The Cajal Body undergoes nucleolar invasion upon SMN depletion. (A) Relative human SMN1 mRNA expression level +/- Dox treatment for 10 days. (B) Mass Spectrometry of protein abundance from NP40 or post-NP40 chromatin extracts +/- Dox for 10 days. (C) Co-IF of TCAB1 and coilin from fixed shSMN cells +/- Dox for indicated days. Scale bars, 5 µm. (D) Co-IF of TOE1 and coilin from fixed shSMN cells +/- Dox for 10 days. Scale bars, 5 µm. (E) Two-channel Confocal live images of shSMN cells stably expressing either mCherry-coilin or GFP-fibrillarin. Four major coilin phenotypes/morphologies were categorized in the image panel on the left. The number of the four categories per cell across the indicated days of Dox treatment was counted and plotted on the right. Scale bars, 5 µm. (F) Live cell holo-tomography images of shSMN treated with Dox for 6 days. RI alone on the bottow row, and the RI-GFP epifluorescence overlay is shown on the top row. T0 indicates an arbitrary time point upon the onset of mitosis. (G) Eccentricity from Coilin-marked compartments, segmented, and quantitated by CellProfiler during the time course of Dox treatment. Kruskal-Wallis test considering the non-normal distributions was used with p≤ 0.0001, n= 27,951. (H) Co-IF of TCAB1 and NPM1 in coilin-proficient and -deficient isogenic cells, +/-DOX for 10 days. Scale bars, 5 µm. (I) TCAB1-NPM1 overlap coefficient from H, at least 300 cells from each conditions were quantitated using CellProfiler. (J) Western blots characterizing the coilin protein level in the isogenic COILIN-CRISPR clones (CKO) derived from the shSMN parental background. (K) Allelic status of the Coilin-KO (H2 clone) by PCR genotyping followed by NGS sequencing. The annotated Coilin ATG is labeled with a green triangle that orients from right to left (5’ to 3’). Total NGS reads for the indicated genotype, and their allelic ratios are shown. Only 2 allelic types have been identified at a roughly 2:1 ratio, consistent with the triploid status at the Coilin locus.

**Fig.S2.**
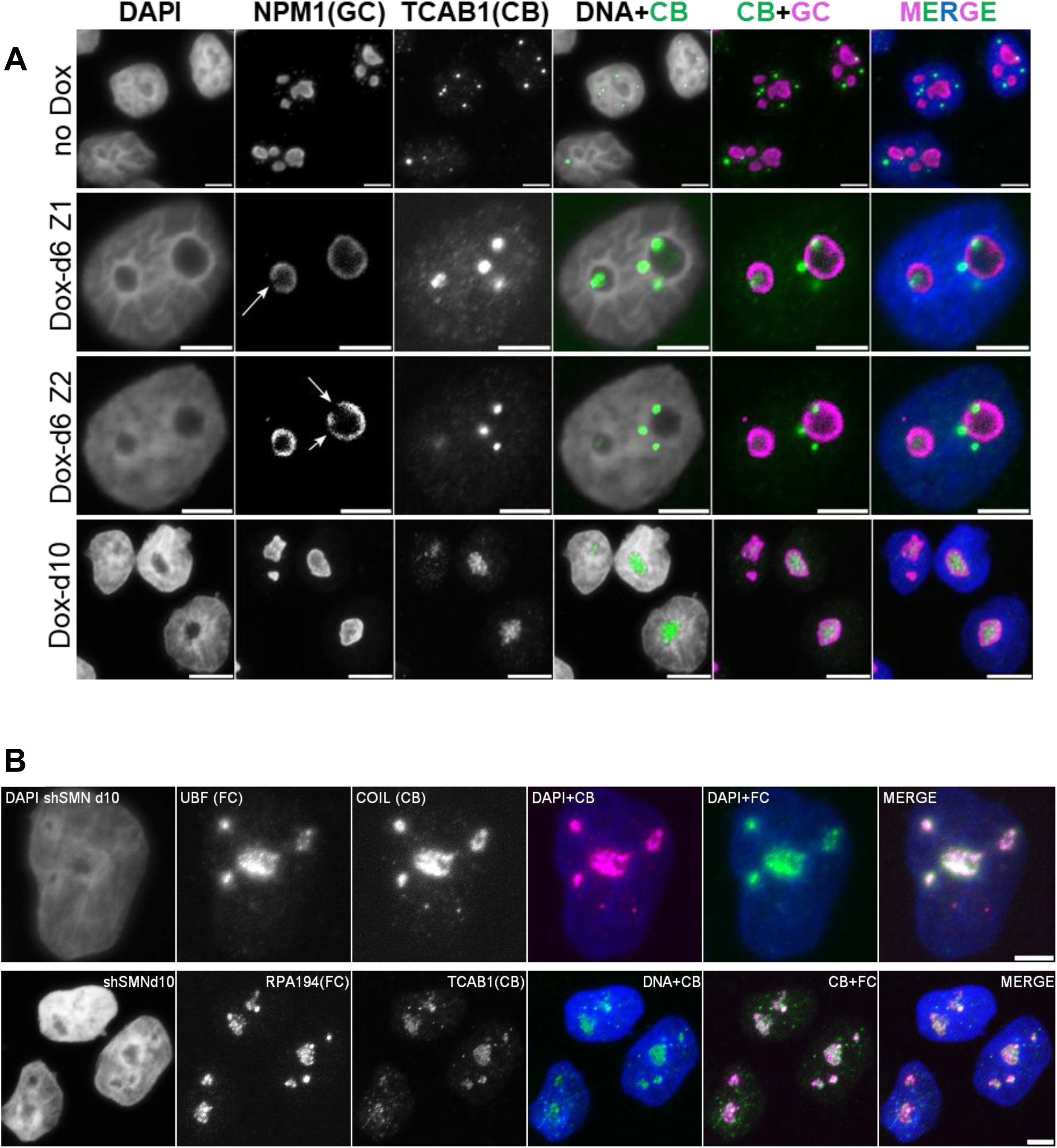
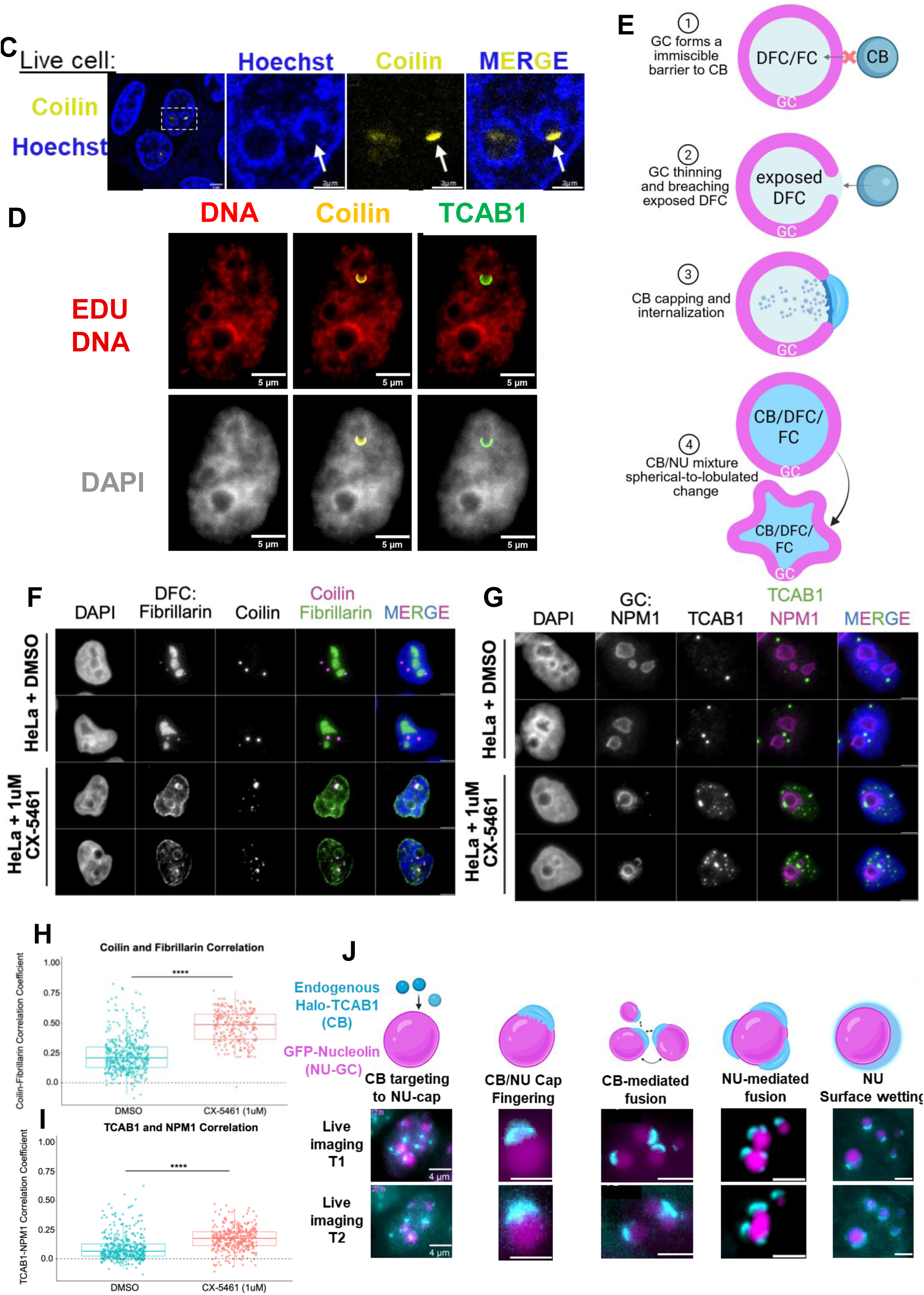
Nucleolar migration of CB involves the breaching of the immiscible GC barrier. (A) Co-IF between NPM1, a GC marker, and TCAB1, a CB marker, upon 6 or 10 day treatment of Dox. Maximum Intensity Projections of Z-stacks are shown except for two single Z slices, Z1 and Z2, on the 6th day of Dox-treatment. Scale bars, 3 µm. (B) Co-IF between CB proteins, coilin and TCAB1, and FC proteins, UBF and RPA194. Scale bars, 2 µm. (C) Confocal live image of 10-day Dox-treated shSMN cells transfected with EGFP-Coilin. EGFP signal in yellow, DNA labelled by Hoechst in blue. (D) Co-IF of 6-day Dox-treated shSMN cells detecting endogenous coilin and TCAB1. EDU labeling was performed, and the discrete nuclear staining (red) marks late-replication regions, including the peri-nucleolar heterochromatin. CB proteins form a cap-like structure on one of the nucleoli that has a discontinuous EDU and DAPI-positive periphery. (E) A model describing the observed steps CBs take to penetrate the GC barrier during the nucleolar internalization. (F) Co-IF between DFC marked by fibrillarin and CB marked by coilin in parental HeLa cells treated with CX-5461 for 24 hours. (G) Co-IF between GC marked by NPM1 and CB marked by TCAB1 in parental HeLa cells treated with CX-5461 for 24 hours. (H) Degree of Coilin and Fibrillarin colocalization upon CX-5461 treatment (DMSO n=427, CX-5461 n=238), p=1.05e-60. (I) Degree of TCAB1 and NPM1 colocalization upon CX-5461 treatment (DMSO n=401, CX-5461 n=345), p=4.38e-31. (J) Live imaging of CX-5461-treated cells expressing endogenously tagged TCAB1 (Cyan) and transfected GFP-Nucleolin (Purple). Two adjacent imaging frames with 1 min interval are shown. A schematic for liquid-like behavior between two phases is shown.

**Fig.S3.**
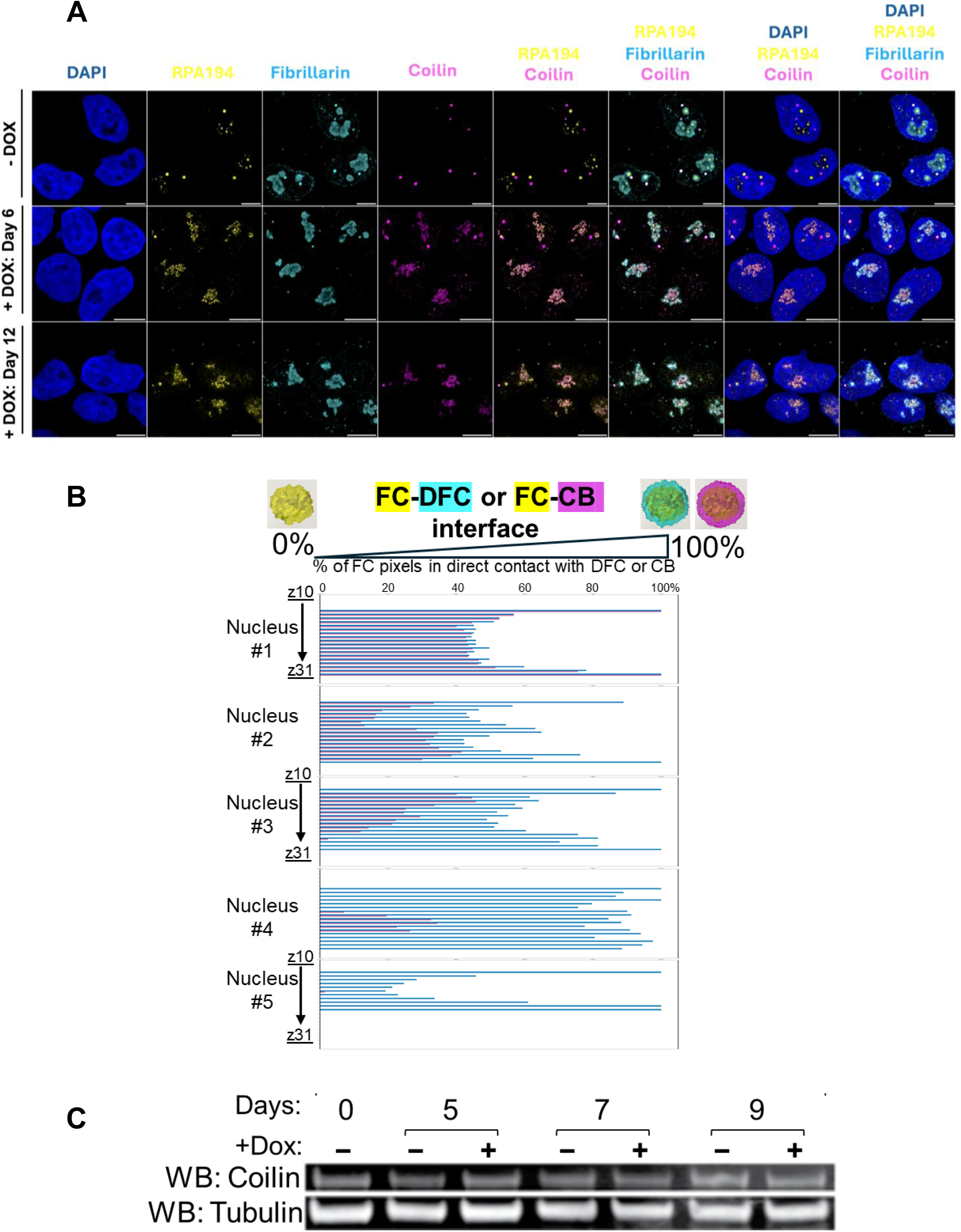
Intranucleolar Coilin binds rDNA at a restricted peri-FC zone. (A) Confocal microscopy of co-IF from HeLa shSMN cells treated with Dox as indicated. FC (RPA194, yellow), CB (Coilin, magenta), and DNA (DAPI, blue). Maximum Intensity Projections of Z-stacks are shown. Last two columns show the zoom-in view of the dotted area on the 4th column. Scale bar, 5 µm. (B) Percentages of FC pixels (yellow) in contact with DFC or CB; all pixels within a single Z-plain are shown. All Z planes from a total of five 3D-reconstructed nuclei (#1-#5) are shown. (C) Western blotting of coilin’s total protein level from a whole cell extract, upon indicated Dox treatment.

**Fig.S4.**
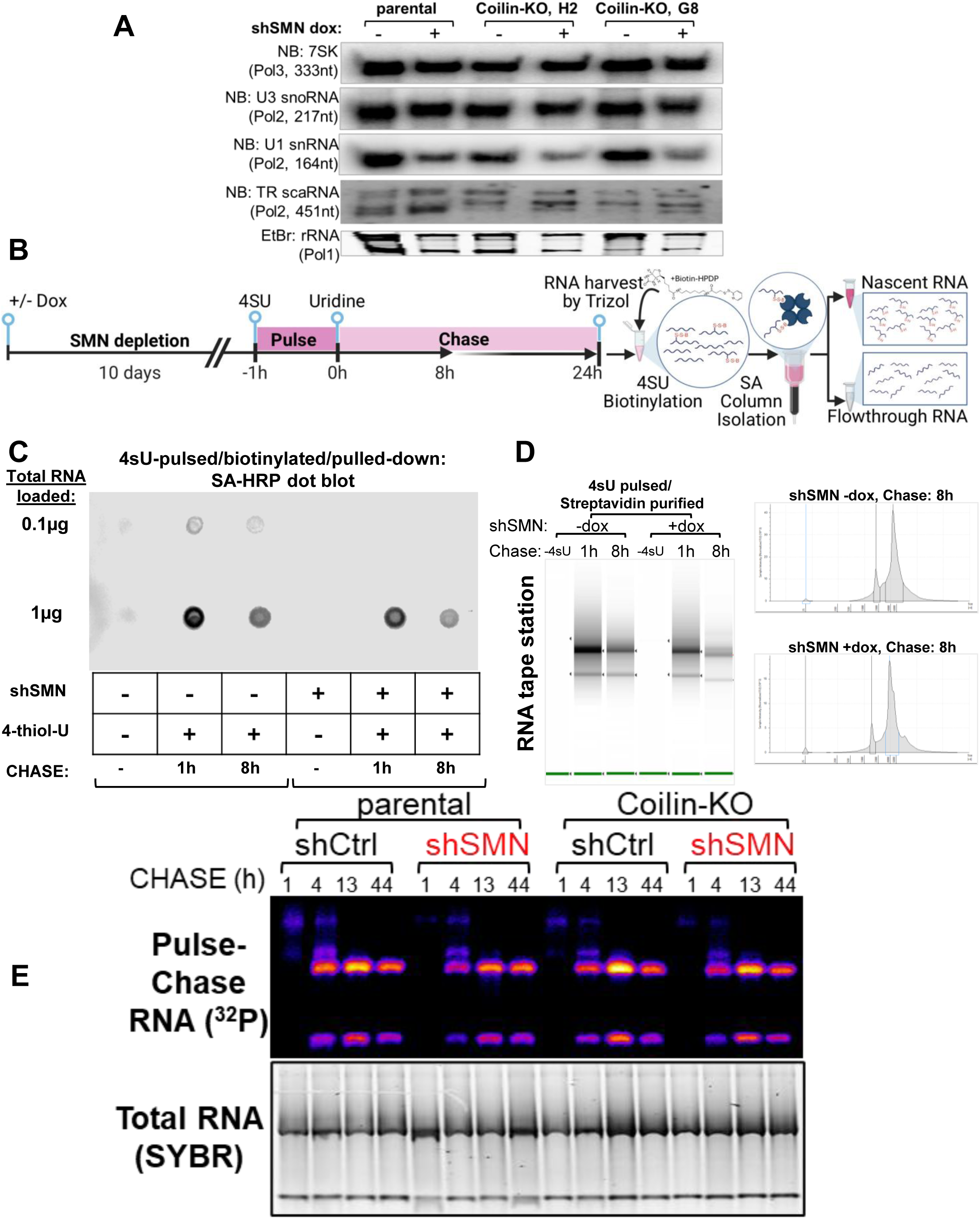
Intranucleolar Coilin impedes rRNA transcription. (A) Northern blotting shows steady state RNA levels from the parental shSMN cell line, and two derivative clones from Coilin CRISPR (H2 and G8). +/- Dox treatment for 10 days. (B) An orthogonal nascent RNA pulse-chase pipeline using 4SU or 4-thio-Uridine, followed by in vitro biotinylation with Biotin-HPDP or N-[6-(Biotinamido)hexyl]-3’-(2’-pyridyldithio)propionamide, and SA or Streptavidin column purification, with SA column-bound RNA fraction as “nascent”, and column flowthrough fraction as “steady state”. (C) Dot blot quantitation of 4SU incorporation efficiency from total RNA (0.1 or 1µg), visualized with SA-HRP or Horseradish Peroxidase-conjugated Streptavidin. 4SU-negative mock control and 4SU-pulsed samples with indicated chase times are included, +/- Dox treatment for 10 days. (D) 4sU pulse-chase nascent RNA fractions, purified by Streptavidin column and visualized by capillary electrophoresis (Tape station). Gel image (left) and electropherograms (right) are shown. (E) Pulse-chase labeling of nascent rRNA production using 32P-orthophosphate (Fig.4E-F), in either shCtrl or shSMN expressing cells, with or without Coilin detected expression, treated with dox for 10 days. (Top) the agarose-MOP gel is exposed by autoradiography to visualize labeled rRNA species at the indicated chase time; (Bottom) the exact same gel was stained with SYBR-Green-2 dye for normalization purposes.

**Fig.S5.**
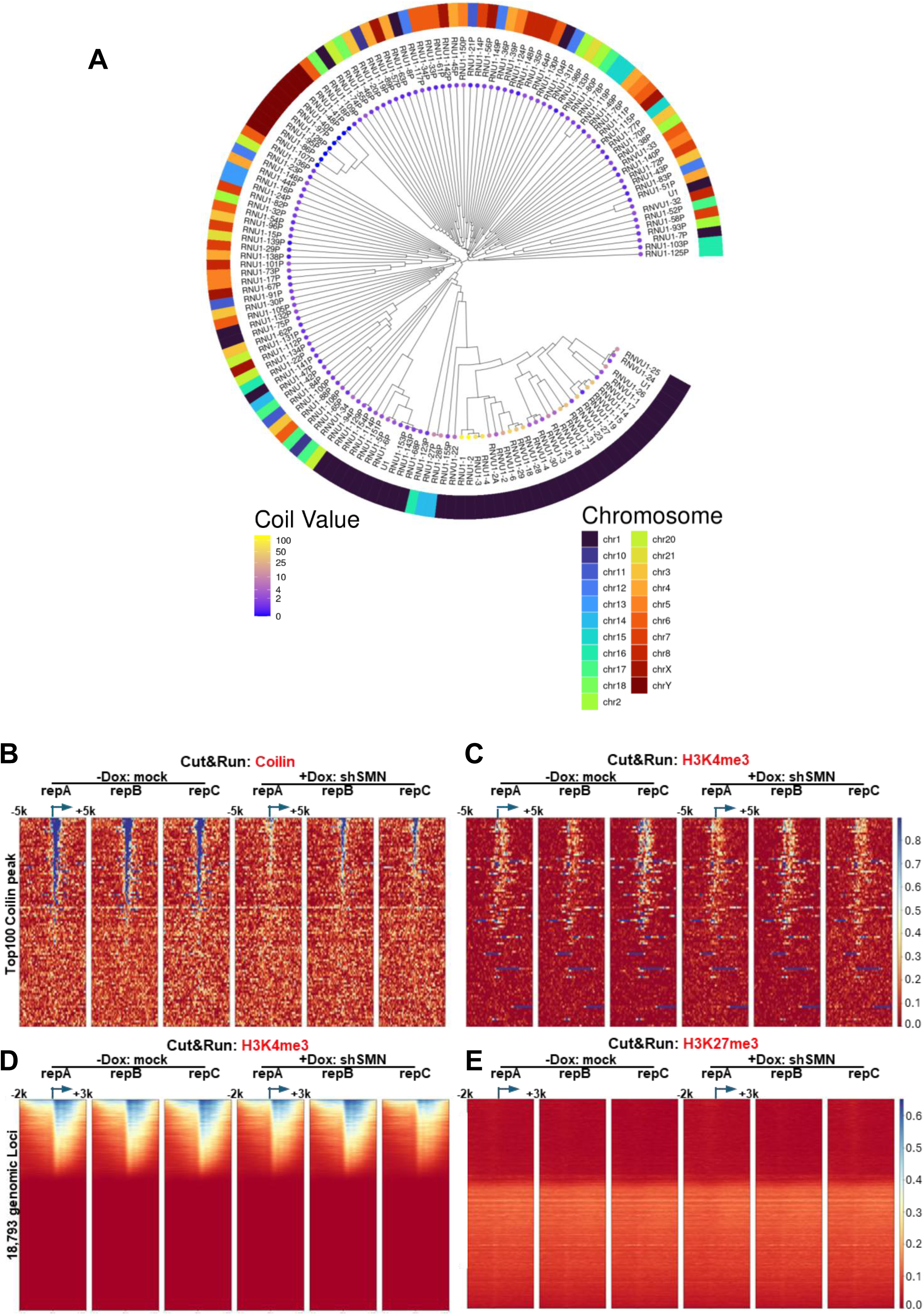
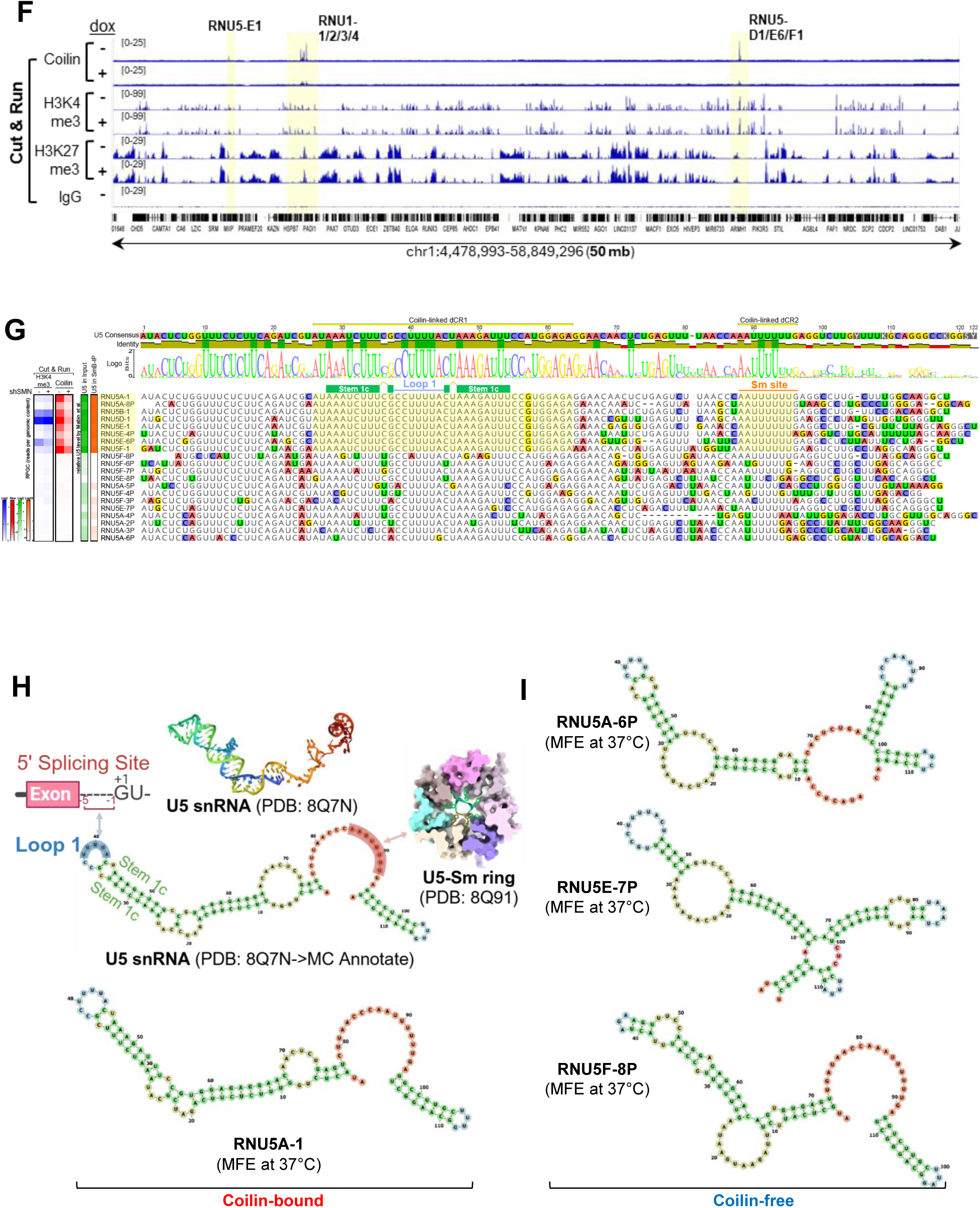
Coilin’s genomic targeting functions post-transcriptionally. (A) Phylogenetic analysis of U1 snRNA gene promoters. A Maximum Likelihood tree was constructed using the 500 nt proximal promoter sequences of annotated U1 genes. (Inner) Branches are colored according to chromosomal origin. (Outer) The heatmap ring displays the relative Coilin CUT&RUN signal intensity for each locus (Purple = low; Yellow = high). (B-E) Heatmap ranking of CUT&RUN signals using anti-coilin, H3K4me3, and H3K27me3 antibodies; 5-10kb genomic window centering around the TSS, which is denoted by an arrow. For A and B, top 100, per coilin signal, of the 4,600 of annotated snRNA loci were ranked; for C and D, all 18,793 gene promoters are rank-listed. (F) IGV view of the U1/U5 snRNA clusters on chromosome 1 over a 50 megabase (mb), with coding regions highlighted in yellow. Local CUT&RUN signals for coilin, H3K4me3, and H3K27me3 were shown with IgG as a negative control. (G)Multiple sequence alignment of U5 snRNAs. CUT&RUN of Coilin and H3K4me3 signal were annotated, together with U5 abundance in either steady-state (Input) or SmB-interacting fraction (SmB-IP). Differentially Conserved Regions, Stem/Loop1 (dCR1) and Sm site (dCR2), are highlighted. (H) CryoEM-derived U5 snRNA structure model, according to PDB: 8Q7N. Stem/Loop1 and Sm site known roles illustrated; secondary structural prediction of RNU5A-1, one of the Coilin-bound U5 snRNA, using its Minimum Free Energy at 37°C. (I) Minimum Free Energy models of U5 snRNA derivatives that are not bound by Coilin.

**Fig.S6.**
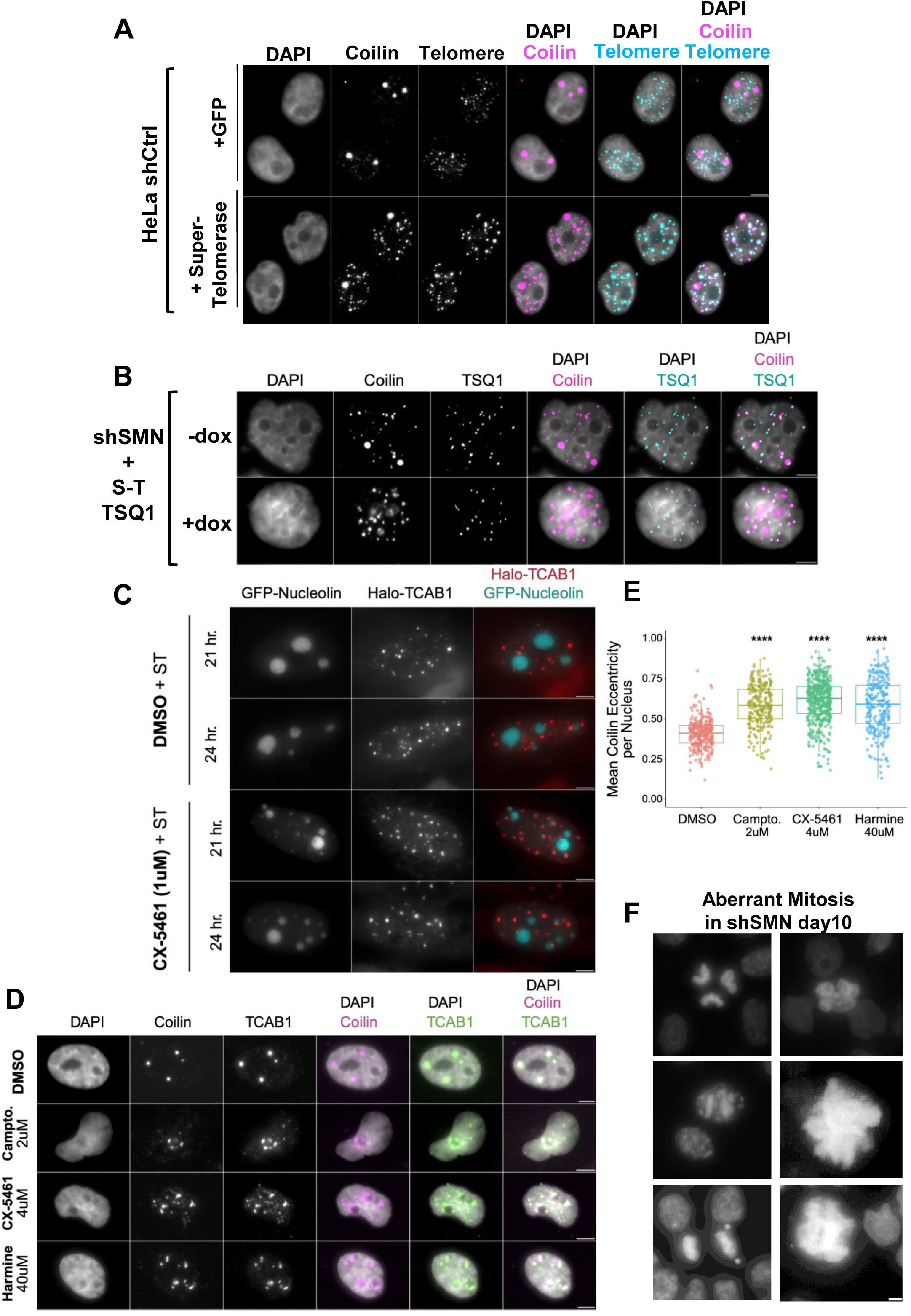
CB-nucleolar mixing impedes Coilin and telomerase scaRNP targeting. (A) ST reconstitution sufficiently targets coilin (Pink, IF) to telomeric foci (Cyan, DNA-FISH). GFP-transfected cells serve as a mock control. Scale bar, 4 µm. (B) ST reconstitution with the TSQ1 variation, +/- Dox treatment for 10-days. De novo telomere elongation was visualized by DNA-FISH using a TSQ1-specific probe. coilin IF shows nucleolar retention signal in the +dox. TSQ1 FISH is quantitated in Fig.6 E-F. Scale bar, 4 µm. (C) Live cell movie frames showing endogenously tagged Halo-TCAB1 HeLa cells transfected with GFP-Nucleolin, +/-CX5461 treatment for the indicated hours. Fragmented Nucleolin structures (Red) are notable in cells treated with CX5461 and are often juxtaposed with TCAB1 (Cyan). Scale bar, 4 µm. (D) Co-IF of the induced nucleolar capping phenotype. Coilin (Purple) and TCAB1 (Green) colocalizing around the DAPI-low structures upon treatment. Scale bar, 4 µm. (E) Coilin eccentricity quantitated from D. (F) Examples of aberrant mitotic cells as quantitated in Fig.6K. Cells were fixed and stained with DAPI to visualize DNAs. Scale bar, 4 µm.

## Notes

### Competing Interest Statement

The authors have declared no competing interest.

